# The fitness landscape of TEM-1 β-lactamase is stratified and inverted by sublethal concentrations of cefotaxime

**DOI:** 10.1101/2022.07.13.499905

**Authors:** Andrew D. Farr, Diego Pesce, Mark P. Zwart, J. Arjan G. M. de Visser

## Abstract

Adaptive evolutionary processes are constrained by the availability of mutations which cause a fitness benefit – a concept that may be illustrated by ‘fitness landscapes’ which map the relationship of genotype space with fitness. Experimentally derived landscapes have demonstrated a predictability to evolution by identifying limited ‘mutational routes’ that evolution by natural selection may take between low and high-fitness genotypes. However, such studies often utilise indirect measures to determine fitness. We estimated the competitive fitness of each mutant relative to all of its single-mutation neighbours to describe the fitness landscape of three mutations in a β-lactamase enzyme at sub-lethal concentrations of the antibiotic cefotaxime in a structured and unstructured environment. We found that in the unstructured environment the antibiotic selected for higher-resistance types – but with an equivalent fitness for subsets of mutants, despite substantial variation in resistance – resulting in a stratified fitness landscape. In contrast, in a structured environment with low antibiotic concentration, antibiotic-susceptible genotypes had a relative fitness advantage, which was associated with antibiotic-induced filamentation. These results cast doubt that highly resistant genotypes have a unique selective advantage in environments with sub-inhibitory concentrations of antibiotics, and demonstrate that direct fitness measures are required for meaningful predictions of the accessibility of evolutionary routes.

**Importance:** The evolution of antibiotic resistant bacterial populations underpins the ongoing antibiotic-resistance crisis. We aim to understand how antibiotic-degrading enzymes can evolve to cause increased resistance, how this process is constrained and whether it can be predictable. To this end we performed competition experiments with a combinatorially-complete set of mutants of a β-lactamase gene subject to sub-inhibitory concentrations of the antibiotic cefotaxime. While some mutants confer their hosts with high resistance to cefotaxime, in competition these mutants do not always confer a selective advantage. Similarly, we identified conditions involving spatial structure where mutations causing high resistance result in a selective disadvantage. Together, this work suggests that the relationship between resistance level and fitness at sub-inhibitory concentrations is complex; predicting the evolution of antibiotic resistance requires knowledge of the conditions that select for resistant genotypes and the selective advantage evolved types have over their predecessors.

## Introduction

The mapping of genotypic space with fitness is a primary concern of evolutionary biology. This interest arose because this mapping bridges genetics and evolution, being shaped by the interactions between mutations and governing which mutational trajectories evolution can follow. Since 1932 (1) the ‘fitness landscape’ metaphor has been used to visualise the complicated mapping of genotype to fitness. More recently, molecular biology has made possible the construction of mutational networks where sets of mutations – in either one (2-8) or multiple genes (9-13) – are combined and expressed in an experimental strain. The interactions among mutations generate the topography of these empirical fitness landscapes, and render particular trajectories across the landscape more likely than others. In a foundational study, Weinreich and colleagues demonstrated that of the 120 possible trajectories linking an ancestral ß-lactamase antibiotic resistance allele with a highly-adapted genotype with five mutations, only 18 trajectories are selectively accessible (3). The level of constraint within such landscapes suggests a certain degree of predictability of evolutionary processes, although the relation between epistatic constraints and evolutionary predictability is complicated by the presence of multiple adaptive peaks (14, 15).

For empirical fitness landscapes to inform evolutionary biology and enable meaningful predictions regarding evolutionary processes, they require informative measures of the relative fitness of each genotype in the landscape. However, the actual measures of fitness used to form such fitness landscapes generally do not represent the outcome of direct competition between the genotypes investigated. In real life, *de novo* variants often directly compete with their progenitors, making it essential to use fitness measures that capture their competitive ability to predict their fates. Despite their name, empirical fitness landscapes are often quantified by the functionality of a focal enzyme. The mutational networks in dihydrofolate reductase (2), methyl-parathion hydrolase (16) and ß-lactamase (3, 5, 8, 17) have used the activity of the enzymes and their effects on organismal growth or survival as proxies for fitness. Many empirical fitness landscapes do provide measures of the relative fitness of genotypes. However, with some notable exceptions (18), these fitness measures are made relative to a common competitor – not immediate evolutionary predecessors – and may be complicated by non-transitive interactions among the genotypes, causing deformations of the landscape due to eco-evolutionary changes of the environment (19). Such environmental alterations may occur if genotypes alter the extracellular concentration of a common resource or inhibitory substance available to co-cultivated genotypes (20-23). Although technically challenging, fitness landscapes would ideally present relative fitness measures of each genotype to mutational neighbours from which they arise.

We constructed an empirical fitness landscape measured by the competitive fitness of each genotype in the landscape relative to its immediate mutational neighbourhood. This landscape consists of combinations of three mutations which underpin the evolution of the ß-lactamase TEM-52, a clinically relevant member of the extended spectrum ß-lactamases (ESBLs). Following the clinical introduction of cephalosporin antibiotics, ESBL’s were selected with the capacity to bind and hydrolyse cephalosporins such as cefotaxime (CTX), resulting in genotypes such as TEM-52. TEM-52 evolved from the ancestral TEM-1 ß-lactamase, which effectively degrades ß-lactams such as ampicillin, but has low activity against cefotaxime (24). The evolution of TEM-52 from TEM-1 involved three non-synonymous mutations, causing amino acid substitutions E104K, M182T and G238S, which in various combinations form an eight-genotype mutational landscape (see Fig. 1A).

**Figure 1:**
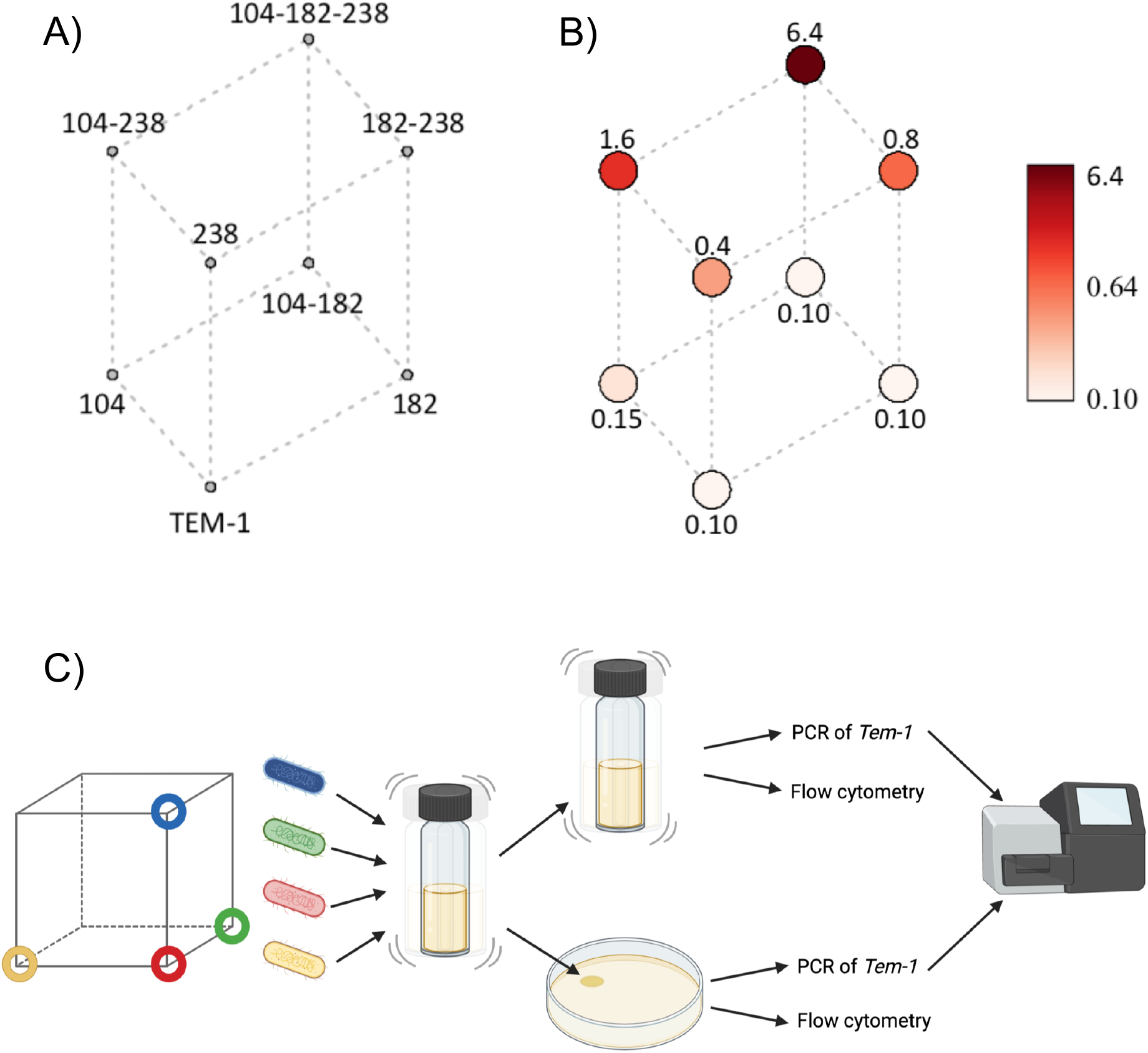
Levels of resistance across the 8-genotype landscape of TEM-1 β-lactamase for cefotaxime resistance and the method used in measuring fitness. A) Cube diagram of the 8-node mutations (numbers refer to amino acid positions of substitutions) with B) red circles overlaid representing median MIC measures in µg/mL of CTX (see table 1). C) Summary of the experimental design to measuring relative fitness of genotypes in the landscape. A focal genotype and the three mutational neighbours with Hamming distance one (each represented by a different colour) were each cultured and mixed together in equal ratios (T0 sample). The mixture of genotypes was then inoculated into either liquid M9 media or spotted onto M9 media solidified with agar either without or with 0.02 or 0.04 µg/mL of CTX. Competition mixtures were sampled at 0, 24 and 48 hours, whereupon cell concentrations were measured by flow cytometry, or samples were stored for amplicon sequencing to determine the ratio of mutations present.

To assess the adaptive benefit of the mutations across the landscape, we reconstructed this set of eight genotypes into a common ancestral strain and competed each genotype with its three neighbouring genotypes with a Hamming distance of one. Competitions were performed in a structured (on the surface of agar-solidified medium) and an unstructured (in liquid medium) environment, to investigate the role of spatial structuring on the fitness of competitors. Spatially structured environments – such as colonies grown on agar surface – were previously shown by Frost and colleagues (25) to provide a selective benefit to non-resistant strains when co-cultured with resistant strains. This selective benefit was due to the induction in sensitive strains of filamentous cellular morphologies – a physiological response of sensitive bacteria exposed to near-inhibitory concentrations of ß-lactam antibiotics (26). These elongated cells helped the non-resistant strains to invade the unoccupied space of the colony edge, where they presumably had better access to resources, ultimately resulting in a selective benefit over non-elongated resistant strains (25). We wanted to see whether this phenomenon was reproducible in our experimental system, and if so, how spatial structuring alters the topography of the fitness landscape. Our fitness assays were performed in sub-inhibitory concentrations of CTX – which we define as the range of antibiotic concentrations which permit monocultures of a genotype to grow. The choice of these low concentrations was driven by two considerations: 1) the practical need to measure fitness requires the presence of competitors after a growth period, and high CTX concentrations may completely eliminate low-resistance competitors, 2) the growing interest in sub-inhibitory concentrations of antibiotic as representative of concentrations in environmental settings such as waste-water and sediment (27), and the potential for such sub-inhibitory concentrations to select for resistant genotypes (13, 28-30). When competed in these sub-inhibitory concentrations, our set of genotypes display fitness landscapes that are highly conditional on both antibiotic concentration and spatial structuring of the environment.

**Table 1:** Genotypes used in this study and associated MIC values. DA28202 was the parent strain for all subsequent TEM-1 mutation used in this study. DA28202 expresses a BFP, allowing cellular counts to be made by flow cytometry. Median MIC assays of the TEM-1 mutants were performed on all four biological replicated used in the landscape fitness assays. The median MIC of DA28200 and DA28202 was measured by visual observation and median MIC of the TEM-1 was performed at a separate occasion using OD600 measures of growth (see methods).

| Strain as referred in text | Genotype | Median MIC | Reference |
| --- | --- | --- | --- |
| DA28200 | E coli MG1655 *galK::SYFP2-FRT* | 0.1 | (29) |
| DA28202 | E coli MG1655 *galK::mTagBFP2-FRT* |  | (29) |
|  | (parent of below strains) | 0.04 |  |
| TEM-1 | *galK :: TEM-1* | 0.1 | This study |
| M182T | *galK :: TEM-1* M182T | 0.1 | This study |
| E104K | *galK :: TEM-1* E104K | 0.15 | This study |
| G238S | *galK :: TEM-1* G238S | 0.4 | This study |
| M182T-E104K | *galK :: TEM-1* M182T E104K | 0.1 | This study |
| M182T-G238S | *galK :: TEM-1* M182T G238S | 0.8 | This study |
| E104K-G238S | *galK :: TEM-1* E104K G238S | 1.6 | This study |
| E104K-M182T-G238S | *galK :: TEM-1* E104K M182T G238S | 6.4 | This study |

## Results

### Experimental model: the TEM-1 to TEM-52 mutational landscape

The genotypes studied here were made by insertion of TEM-1 alleles, reconstructed with all combinations of the three mutations that comprise TEM-52, into the chromosomal *galK* locus of *Escherichia coli* MG1655 expressing blue fluorescent protein (BFP) (see Table 1 and the Methods section). To compare the enzymatic effects of the resulting mutant enzyme with fitness, we performed minimum inhibitory concentration (MIC) assays with each biological replicate used in the fitness assays (see methods, Table 1 and Fig 1B). MIC assays were conducted over a 24-hour period starting with a low initial cell density (∼ 1.5 × 10^5^ cells mL^-1^), using the same liquid media used in fitness assays (see Table 1 for results). As expected, the magnitude of these resistance measures was consistently lower than previously measured for plasmid-encoded TEM-alleles (3), likely due to gene dosage arising from the single copy of this gene in the chromosome. In keeping with other studies on the enzymatic consequences of these mutations (3, 5), particular combinations of mutations caused a much higher level of resistance than others (see Fig 1b). Of the single mutations, the G238S amino acid change causes a large increase in resistance, and similarly, E104K-G238S is the most resistant of the double mutants. The E104K-M182T-G238S triple mutant confers the highest level of resistance of this landscape of mutants.

### Paired competitions describe inverted and stratified resistance-fitness relationships depending on the environment

To understand the effect these mutations may have on relative fitness of each strain at low antibiotic concentrations, we initially performed pairwise-competitions. Each of the eight genotypes of the landscape was competed with a common reference strain labelled with chromosomally-encoded yellow fluorescent protein (YFP) which did not express TEM-1 (and hence did not contribute to hydrolysis of CTX). Fitness assays were performed in structured (agar media) and unstructured (liquid media) environments containing 0, 0.005, 0.01, 0.02, 0.04 and 0.08 µg/mL of CTX, and cell numbers were measured by flow cytometry (see methods). Pairwise fitness assays of these strains in the absence of CTX revealed no significant relationship between the alleles’ ranked resistance and relative fitness, hence the alleles showed no trade-off between resistance and competitive fitness (see Fig. 2 and supp. Fig S1A and B). Not surprisingly, expression of higher resistance alleles was beneficial in the unstructured environments supplemented with 0.005 µg/mL to 0.08 µg/mL of CTX (see supp. Fig. S1A). This observation was pronounced for all genotypes featuring the G238S mutation. At both 0.02 µg/mL and 0.04 µg/mL, all genotypes which include the G238S mutation had a similar mean selection coefficient, despite an approximate 16-fold difference in the median MIC of genotypes G238S and E104K-M182T-G238S (see Table 1). The absence of a selective benefit between these highly resistant types was confirmed in competitions with multiple genotypes (see below).

**Figure 2:**
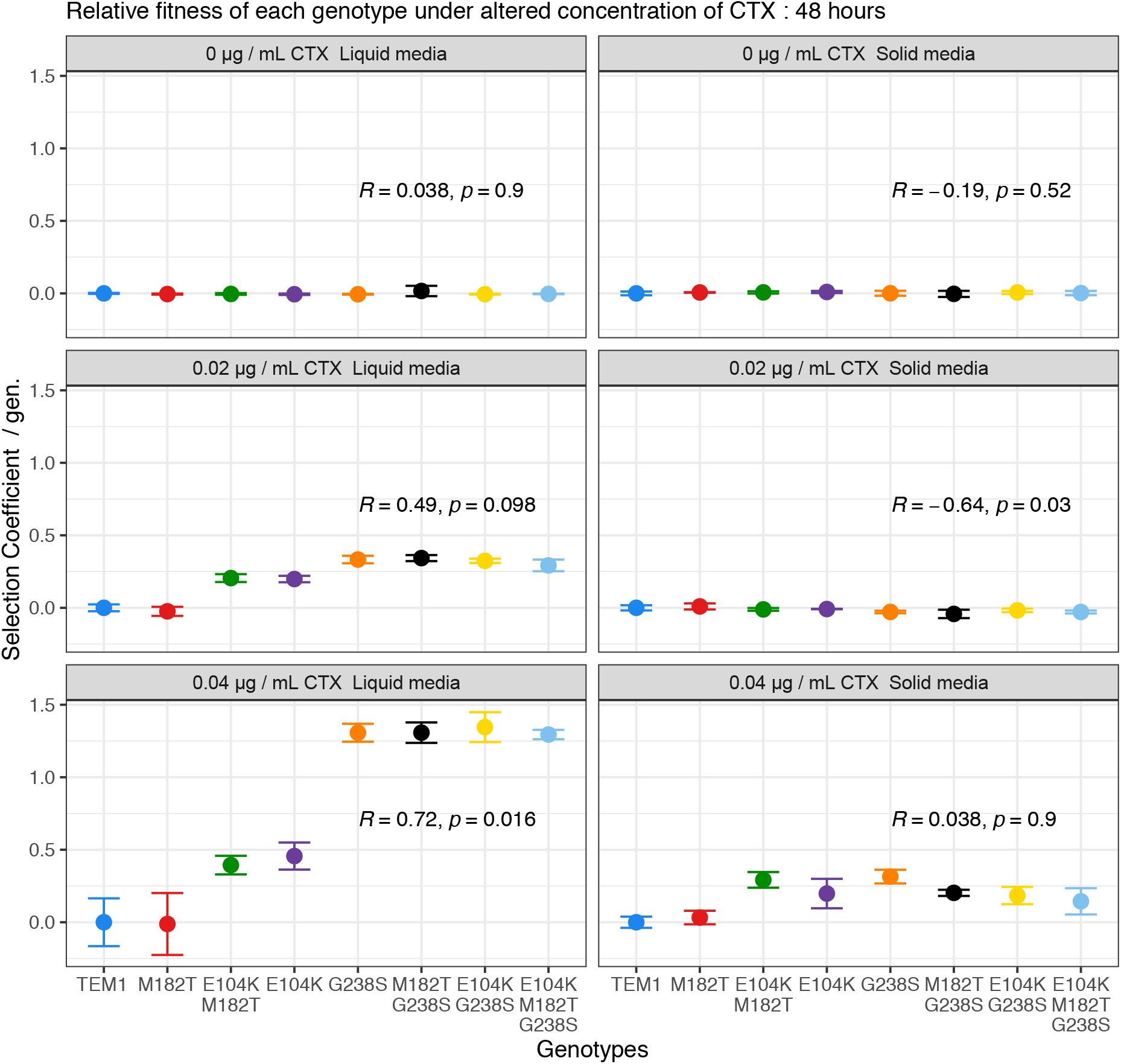
Pair-wise fitness assays relative to MG1655-YFP. Relative to a common competitor, the individual genotypes (ordered by MIC values, ranked from lowest (TEM-1) to highest (E104K-M182T-G238S) exhibited different competitive advantages in the presence and absence of spatial structuring. In liquid media, more resistance types exhibited higher selection coefficient values, which was significant at 0.04 µg/mL of CTX. Competed on solid media, higher resistant types had significantly lower selection coefficient values. Data points represent the means of three biological replicates, error bars represent standard deviations and statistical tests presented are Kendall rank correlation coefficient Rho and p-values.

In structured environments, the relationship between the resistance of each genotype and their relative fitness in pairwise competitions was more complex. Fitness differences among genotypes were much smaller than in liquid cultures (Fig. 2). After both 24 and 48 hours of competition in 0.005, 0.01 and 0.02 µg/mL of CTX, we found a significantly negative correlation between resistance of the genotypes and their fitness (see supp. Fig. S1B). Assays performed at 0.04 µg/mL no longer showed a negative relationship between resistance and fitness, but no significant positive relationship was observed after either 24 or 48 hours of competition (Fig. 2). The highest concentration of 0.08 µg/mL of CTX, resulted in low or no recovery of low resistance competitors. Based upon this observation of the low recovery of the less-resistant genotypes at 0.08 µg/mL of CTX, further multiple-genotype competitions (see below) were conducted at lower concentrations of 0.02 and 0.04 µg/mL CTX.

### Relative fitness of each genotype relative to its three single-mutation neighbours

We wanted to test the impact of selection with antibiotics across the whole fitness landscape, and to make direct measures of fitness between neighbouring genotypes of the 3-mutation fitness landscape. To do so, we competed each of the eight genotypes with the three neighbouring genotypes, resulting in eight competitions involving four competitors (see methods for details and supp. Fig S1). We performed fitness assays at three concentrations of CTX – at the concentration where sensitive genotypes had a growth disadvantage in pairwise comparisons, but were still recoverable (0.04 µg mL^-1^), at half this concentration (0.02 µg mL^-1^) and in environments without CTX. Using these concentrations, we could observe the effect of selection by CTX on the growth of all strains of the landscape. We utilised amplicon sequencing to determine the relative fitness of four competitors simultaneously. Deep sequencing data were analysed with an approach that is insensitive to the effects of recombination between TEM alleles, caused by low-rate template switching during PCR amplification (see Methods section). In order to derive selection coefficients, we measured the total inoculum size and final yield using flow cytometry (see methods and Fig. 1C).

The relative fitness of particular genotypes in liquid culture is partially predictable by their resistance measures. Measures of fitness were calculated as selection coefficients (see Methods). At both concentrations of CTX in liquid cultures, expression of the G238S mutation afforded genotypes a clear fitness advantage relative to non-G238S expressing genotypes (see Fig 3 and supp. Fig S2). This general relationship between resistance and fitness is shown by a strongly positive correlation between measures of resistance and fitness of each genotype relative to its three competitors in liquid cultures with 0.02 and 0.04 µg CTX mL^-1^ (see Fig 4 and supp. Fig S3). However, this positive correlation of resistance and fitness was not always apparent. At both concentrations of CTX in liquid culture, the four high resistance genotypes containing mutation G238S had a similar relative fitness (see Fig 4 and supp. Fig S3). To confirm that these four genotypes have a similar fitness benefit in the tested conditions, direct selection coefficient values were calculated for each of these types relative the other G238S-expressing competitors. Despite substantial differences in resistance between the genotypes, there was no significant relationship between resistance and selection coefficient values of these high-resistance genotypes under any tested condition at either time point (see supp. Fig S4).

**Figure 3:**
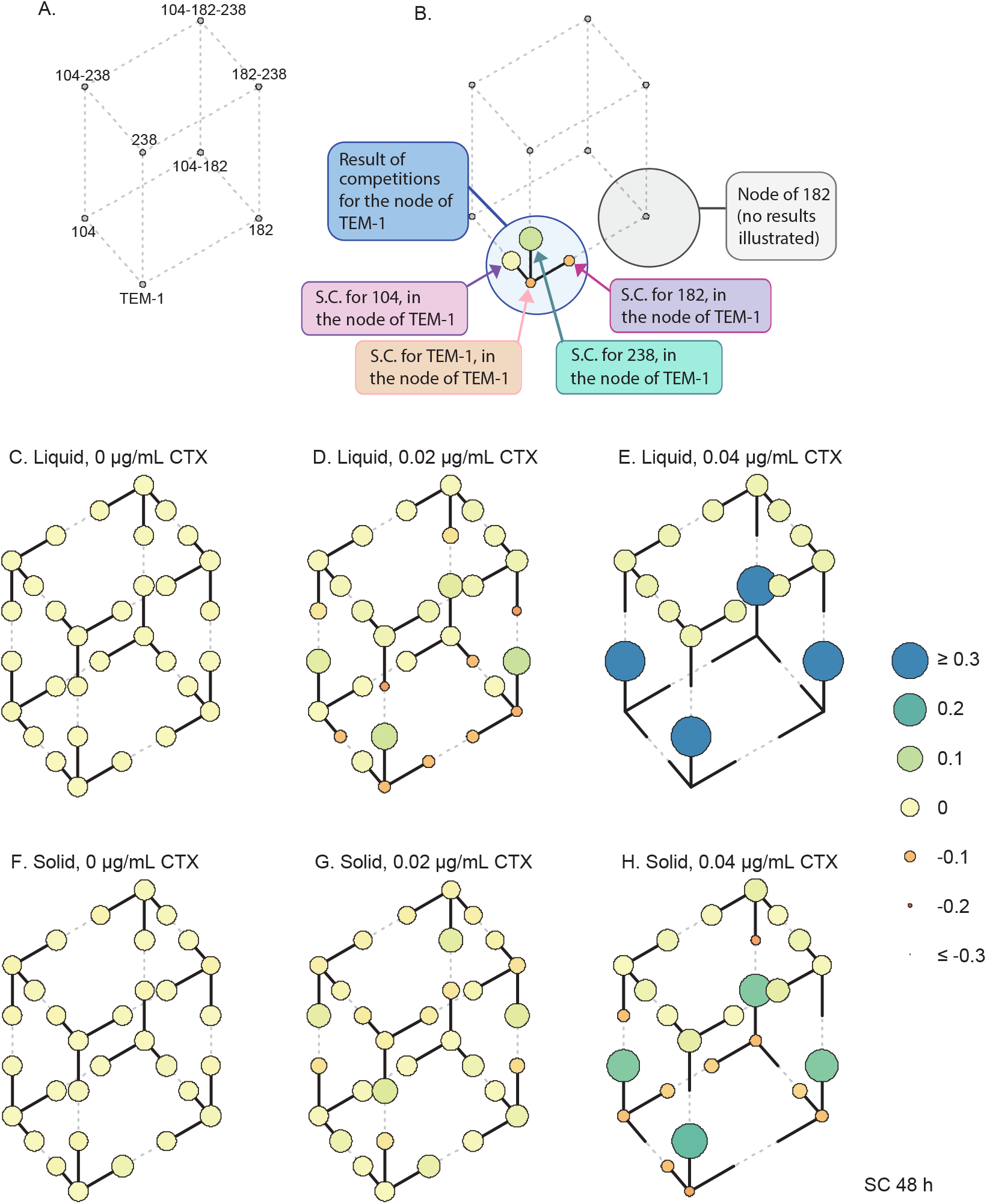
Selection coefficients (SC) of each genotype relative to three genetically-related competitors. **A**. The network of eight genotypes in this study. **B**. Guide illustrating the results of one competition for the node TEM-1. A total of eight competitions were performed – one competition per node – each involving a focal strain, and the three competitors distinct from the ‘focal’ strain by a single mutation. For instance, the node of TEM-1 was comprised of a competition of the focal strain TEM-1, together with E104K, M182T and G238S. **C**. Selection coefficient data of 8 genotypes, each measured relative to three competitors at 48 hours. Competitions were performed in either liquid (C – E) or solid media (F – H) with either 0, 0.02 or 0.04 µg/mL of CTX. SC values are represented by the size and colour of the circles (see legend), and are capped to −0.3 and +0.3 to allow visualization of fitness differences at lower CTX concentrations. Each circle represents the mean SC value of four independent replicates.

**Figure 4:**
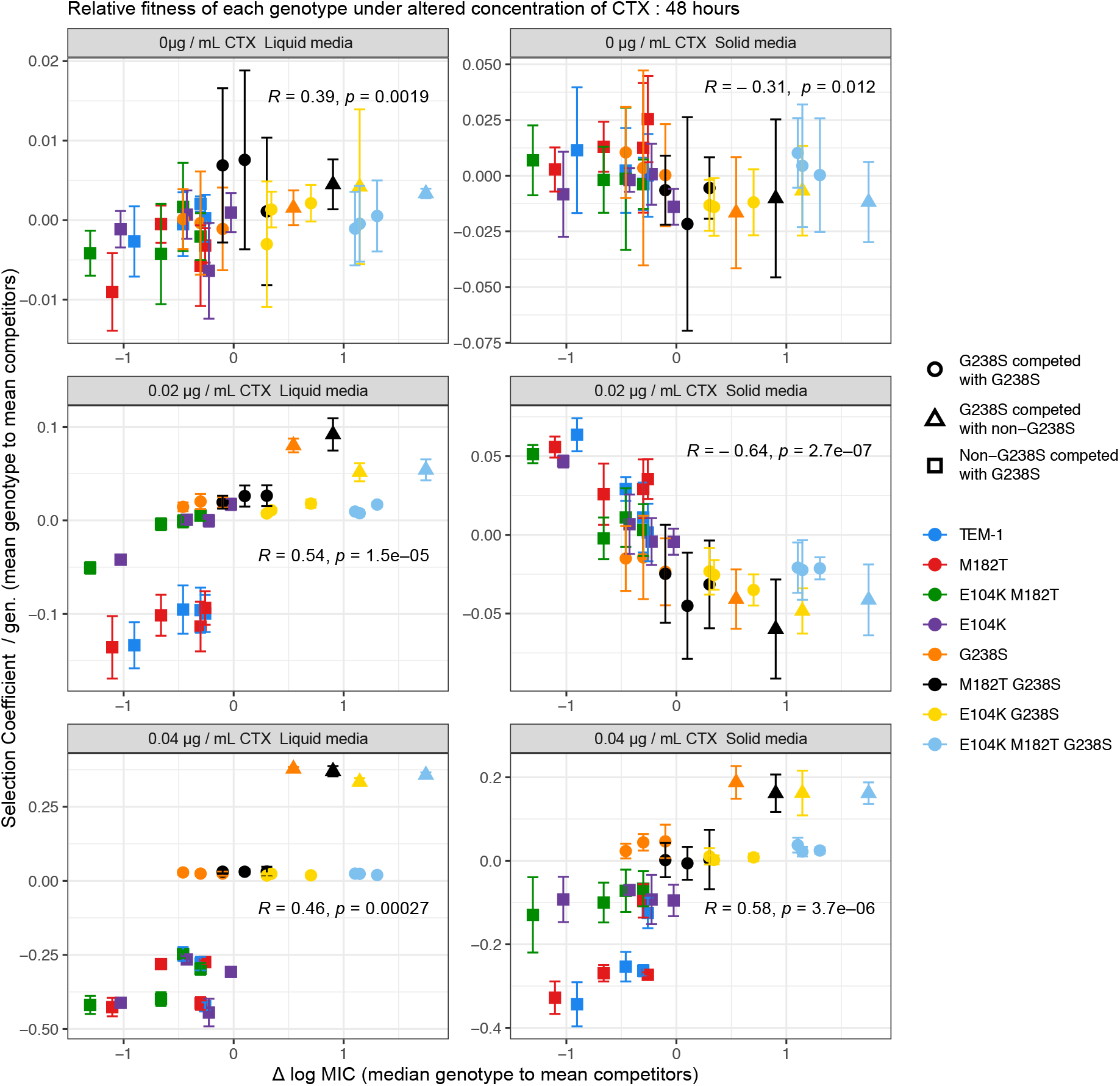
Relationship between resistance and selection coefficient of genotypes competed in 4-genotype competitions over 48 hours. Data points represent the mean selection coefficient of each genotype relative to the three other competitors, and the difference between the log MIC of each genotype and the mean log MIC of the three competitors. A correlation of fitness with resistance is observed at both concentrations of CTX in liquid media, however, little difference is observed between competitors which feature a G238S despite differences in resistance. A similar relationship of resistance and fitness is observed at higher concentrations of CTX for competitions competed on solid media. However, a significantly negative correlation is observed when genotypes were competed on solid media at the lower concentration of CTX. See supplementary figure S2 for similar data at 24 hours. Data points represent the means of four biological replicates, error bars represent standard deviations and statistical tests presented are Kendall rank correlation coefficient Rho and p-values.

Competitions in spatially structured environments resulted in a positive correlation of relative fitness and resistance at the higher CTX concentration of 0.04 µg mL^-1^ CTX (see Fig 4). Similar to the liquid competitions, genotypes expressing the G238S mutation – regardless of additional mutations and level of resistance – had a similar fitness when directly competed with each other, although stratification was somewhat less stringent (see supp. Fig S4). As seen in the pair-wise competitions (Fig 2) – there was a significant negative correlation between fitness and resistance of each genotype in structured environments with 0.02 µg mL^-1^ CTX (see Fig 4). Under these structured conditions with a low CTX concentration, the mixing of high and low resistant types results in selection for low resistance genotypes.

The observation of high relative fitness of low-resistance genotypes competed in a structured environment with a sub-inhibitory concentration of CTX prompted explanation. Previous work has identified filamentation of low resistance types as capable of providing an advantage in direct competitions (25). Flow cytometry measures were used to identify whether our fitness assay conditions were also capable of inducing filamentous cells. To measure the potential of filamentation, each genotype was grown alone in spatially structured and unstructured media with 0, 0.02 and 0.04 µg mL^-1^ of CTX. The degree of filamentation of these monocultures was estimated by observing the distribution of flow cytometry FSC-A measures (see Fig. 5A) – with larger measures suggesting longer cells in the population (31). We found three interesting patterns when analysing these data (Fig. 5B) with a general linear model (Table S1). First, environmental structure affects the fractions of large FSC-A events (*P* < 0.001), indicating a higher prevalence of filaments in liquid medium than in solid medium. Second, there is a two-way interaction between ranked resistance and CTX concentration in their effect on filamentation (*P* < 10^−4^), indicating that less resistant types tend to filament more at higher CTX concentrations. Third, there is a highly significant three-way interaction between ranked resistance, environmental structure and time (*P <* 10^−9^), capturing the decrease in filamentation on solid medium over time, which only occurs for low resistant types (Fig. 5B). These results demonstrate that while cellular filamentation is a predictor of low relative fitness in the presence of antibiotics in monoculture, in the context of mixed genotypes grown as colonies at sub-inhibitory concentrations of CTX, filamentation provides a mechanism by which less resistant genotypes hold a fitness advantage over higher resistance genotypes.

**Figure 5:**
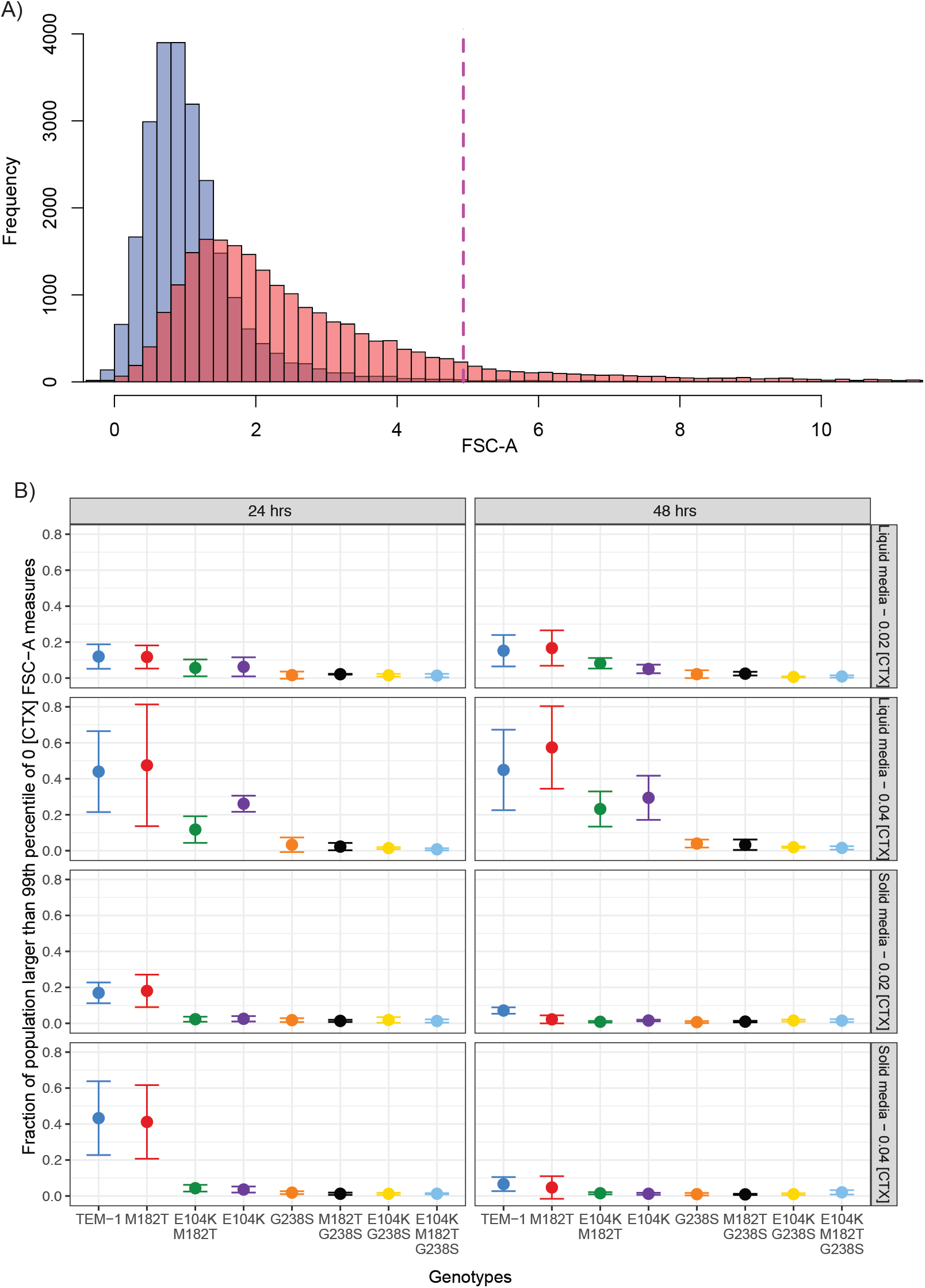
Monocultures of genotypes sensitive to CTX produce larger cells as measured by flow cytometry. A) Representative histogram of the forward scatter (FSC-A, indicating cellular size) of a single replicate TEM-1 genotype grown in liquid supplemented with 0 (blue) or 0.02 µg/mL of CTX (red). The dashed red line indicated the upper 99^th^ percentile of cells grown without CTX as counted in the FSC channel. B) The mean fraction of cells for each genotype grown in monoculture on solid and liquid media supplemented with CTX, exhibiting FSC values larger than the dashed red line in A). Measures of cells were taken after 24 and 48 hours, with the fraction of filaments declining between 24 and 48 hours in populations of less resistant types (TEM-1 and M182T). Data points represent means of three biological replicates, error bars represent standard deviations.

### The large-effect G238S mutation determines fitness and dictates evolutionary fate

Next, we used simple regression models that consider how well fitness can be predicted by (i) CTX resistance, or (ii) the presence of individual beneficial mutations. To consider the relationship between CTX resistance and fitness, we formulated Models 1a and 1b. Model 1a assumes a linear relationship between ΔMIC – the difference in MIC between a genotype and the mean MIC for its node in the landscape – and relative fitness, with a coefficient that scales the relationship. Model 1b limits the range over which the response between ΔMIC and fitness is linear, to capture the effect that beyond a given level of resistance, there is no further effect on fitness. Next to the coefficient to scale the MIC-fitness relationship, this model has a minimum and maximum value of fitness. As an alternative approach, Model 2 divides the genotypes into two fitness classes based on the presence of any one mutation (E104K, M182T or G238S), and assigns each class a fitness value. These models were fitted to the relevant data (Models 1a and 1b: ΔMIC and selection coefficients; Model 2: mutation occurrence and selection coefficients) for each experimental condition separately. Model selection with the Akaike information criterion (AIC) revealed that Model 2, with fitness classes based on the presence of the G238S mutation, provided the best predictions over all conditions (Table 2, Table S2). Support for the different models was comparable in the absence of antibiotics, as the fitness differences between genotypes are minimal. In the presence of CTX, Model 2 was better supported than the other models for 7 out of 8 conditions (Table 2). Estimated model parameters (Table S2) also confirm the main trends observed earlier, including a lower fitness of the G238S-carrying variants in structured media at intermediate antibiotic concentrations (0.02 µg mL^-1^ CTX).

**Table 2:** Summary of model selection results. The Akaike Weight, the likelihood that a model is the best supported model within the set of models tested, is given for each model. CTX indicates the antibiotic concentration (μg/mL) used in the experiment. For model 2, the subscripts indicate the site within the TEM gene used to classify the genotypes. While no model enjoys appreciably higher support in the absence of antibiotics, Model 2_238_ is the best supported model for 7 out of 8 conditions with antibiotics. Complete date for the model selection are given in Table S2.

| CTX | Medium | Time | Akaike Weight |  |  |  |  |
| --- | --- | --- | --- | --- | --- | --- | --- |
|  |  |  | Model 1a | Model 1b | Model 2<sub>104</sub> | Model 2<sub>182</sub> | Model 2<sub>238</sub> |
| 0 | Liquid | 24 | 0.358 | 0.068 | 0.229 | 0.191 | 0.154 |
|  |  | 48 | 0.424 | 0.100 | 0.002 | 0.002 | 0.472 |
|  | Solid | 24 | 0.238 | 0.050 | 0.164 | 0.087 | 0.460 |
|  |  | 48 | 0.190 | 0.551 | 0.038 | 0.029 | 0.193 |
| 0.02 | Liquid | 24 | 0.019 | 0.113 | 0.000 | 0.000 | 0.868 |
|  |  | 48 | 0.019 | 0.108 | 0.000 | 0.000 | 0.874 |
|  | Solid | 24 | 0.227 | 0.528 | 0.000 | 0.000 | 0.245 |
|  |  | 48 | 0.118 | 0.169 | 0.000 | 0.000 | 0.713 |
| 0.04 | Liquid | 24 | 0.000 | 0.000 | 0.000 | 0.000 | 1.000 |
|  |  | 48 | 0.000 | 0.000 | 0.000 | 0.000 | 1.000 |
|  | Solid | 24 | 0.000 | 0.010 | 0.000 | 0.000 | 0.990 |
|  |  | 48 | 0.000 | 0.011 | 0.000 | 0.000 | 0.988 |

Finally, we used the competitive fitness data to simulate evolution on TEM-52’s fitness landscape, with the goal of predicting evolutionary endpoints. We kept the simulations as close to empirical data as possible, predicting what would happen if we would follow our experimental setup over multiple rounds of passaging. At the start of each passage, the seeding genotype and its three single-mutation neighbours are present at equal frequencies. We used an estimate of the number of generations within a passage and the selection coefficients to predict the final frequencies of genotypes, and then randomly selected a single individual to seed the next round of passaging. TEM-1 was assumed as the starting point for each simulation. As a contrast to our experimental data, we generated a prediction for conditions in which resistance is the predominant determinant of fitness, based on the predicted fitness values from Model 1A for liquid medium with 0.04 µg mL^-1^ of CTX (Table S2). Under these conditions, the high-resistance triple mutant TEM-52 predominates (Fig. 6A). Predictions based on our competitive fitness data show different trends. First, in the absence of CTX, all evolutionary endpoints are equally likely, as there are no appreciable differences in fitness (Fig. 6B, 6E). Second, in the presence of CTX the evolutionary endpoint depended almost entirely on the G238S mutation: all variants with this mutation were likely to be endpoints (Fig. 6C, 6D and 6G), except for structured media with 0.02 µg mL^-1^ of CTX, when all variants that did not carry the G238S mutation were likely to be endpoints (Fig. 6F). Overall, selection of simple models predicting fitness based on our measurements (Table 2) and simulating multiple passages of evolution under our experimental conditions (Fig. 6) confirm the trends we have noted from a first inspection of the data. These results stress the importance of the G238S mutation as the key determinant of fitness in the presence of CTX, and highlight that this mutation stratifies the landscape into low and high fitness variants, which may reverse at low antibiotic concentrations in a structured environment.

**Figure 6:**
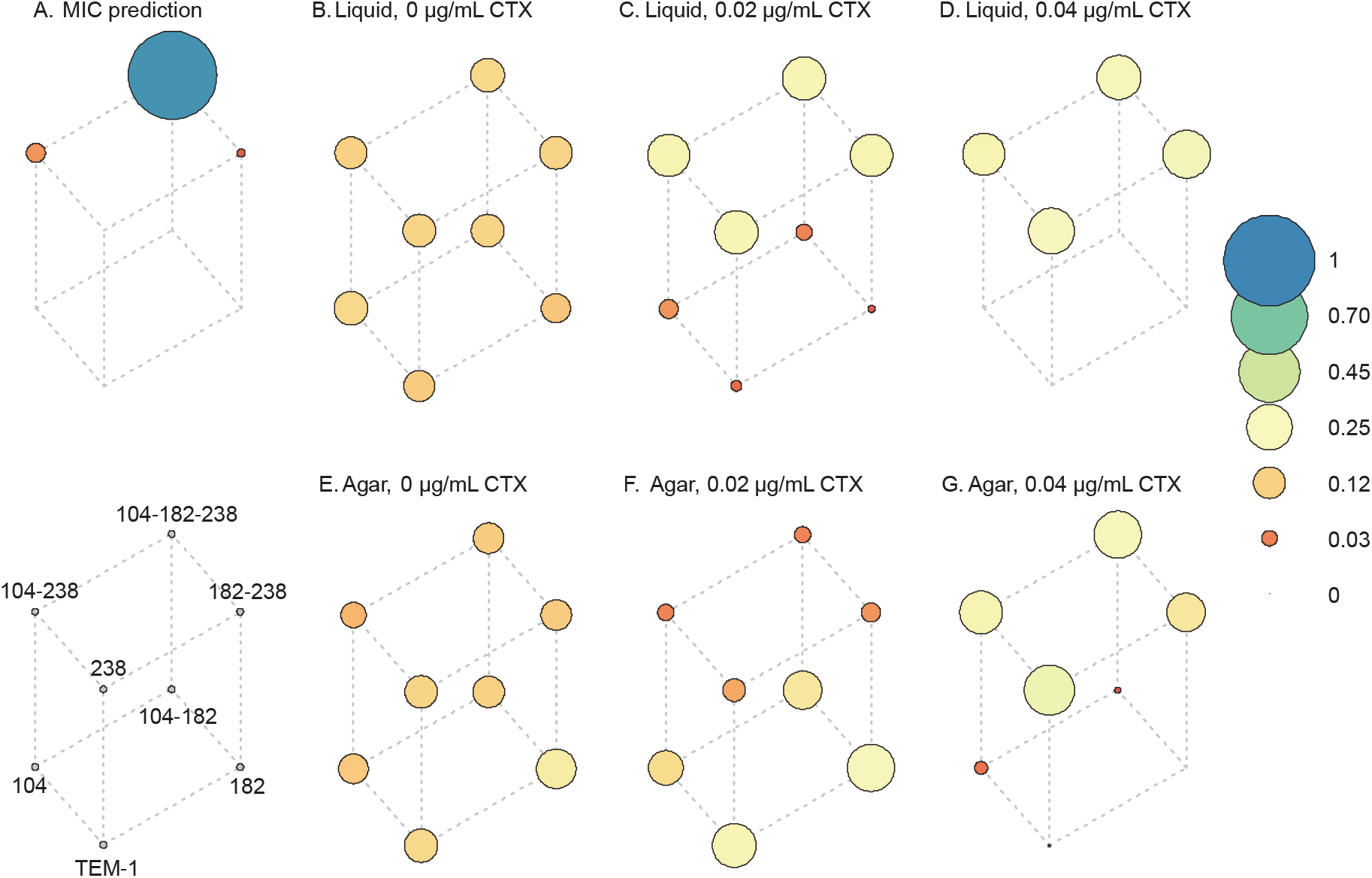
Simulation of evolution by serial passages. The cube represents different genotypes on the three-mutation TEM landscape, with genotype noted in the legend in the bottom left. The circle size and color represent the probability that a genotype will be the final genotype selected after 100 passages of 48 h duration, as indicated by the legend on the far right. These simulations capture conditions identical to our setup for the competition experiments, but running over multiple passages instead of single passage. Each passage is initiated by the starting genotype and the three Hamming distance = 1 mutants at equal frequencies, and the empirical estimates of selection coefficients are used to predict population composition at 48 h. A single allele is randomly drawn and used to seed the next passage as a starting genotype.

## Discussion

Here we report a new approach for investigating empirical fitness landscapes: assaying the fitness of each genotype in the landscape by direct competitions with its local mutational neighbourhood. We think this approach is a useful experimental refinement in the measurement of fitness landscapes, especially in the simulation of circumstances of strong clonal interference (32, 33) or non-transitive fitness interactions. Non-transitive interactions between genotypes (19) could be expected if the ß-lactamase mutants reduced the concentration of CTX, especially in spatially structured environments with lower CTX concentrations. While the approach we used should readily identify non-transitive interactions that lead to the deformation of the landscape – which could be expected if the ß-lactamase mutants reduced the concentration of CTX, especially in spatially structured environments with lower CTX concentrations – no such interactions were observed. Instead, regardless of the number of competitors, fitness was remarkably binary in the presence of antibiotics: genotypes had either high or low fitness, depending on the presence of large-effect mutation G238S. This strong dependence on a single mutation and its resulting stratification of the landscape, may explain why interactions that are more complex remained undetected. We expect that for landscapes involving a stronger influence of different genotypes on the competitive conditions (e.g. environmental antibiotic concentration) – such as might be seen if this landscape was expressed via a multicopy plasmid with stronger gene dosage – such interactions do exist and can be unveiled by using similar approaches.

The fitness landscapes we measured did not show a single-genotype adaptive “peak”, but rather a “stratification” of genotypes with equivalent fitness under the tested conditions. In both structured and un-structured environments, the high-resistance genotypes in our experimental landscape had similar fitness at sub-inhibitory inhibitory concentrations of CTX. This similarity in fitness was largely determined by the presence of the mutation with largest effect on CTX resistance, G238S (34). Simulations indicated that any TEM-mutant carrying G238S presents a similarly likely evolutionary endpoint for the evolution of CTX resistance in our model landscape. We suggest this similarity in fitness amongst high-resistance types is due to similar periplasmic concentrations of CTX (where it binds to its target, i.e. penicillin binding proteins) due to faster CTX hydrolysis rates than CTX diffusion rates into the periplasm (see Fig. 7) (35). At increasing concentrations of CTX, low resistant types will not be able to reduce periplasmic CTX concentrations to sub-inhibitory concentrations and will consequently be inhibited. Competing types with high resistance will all be able to render the effective concentration of CTX at the periplasmic target to near zero. Assuming a similar relative fitness of these genotypes in the absence of CTX, the relative fitness of high resistance types will therefore also be similar under sub-MIC CTX concentrations. As the concentration of CTX further increases, the number of high-resistance and fitness-equivalent mutants will decrease until only those that degrade periplasmic CTX to sufficiently low concentrations survive.

**Figure 7:**
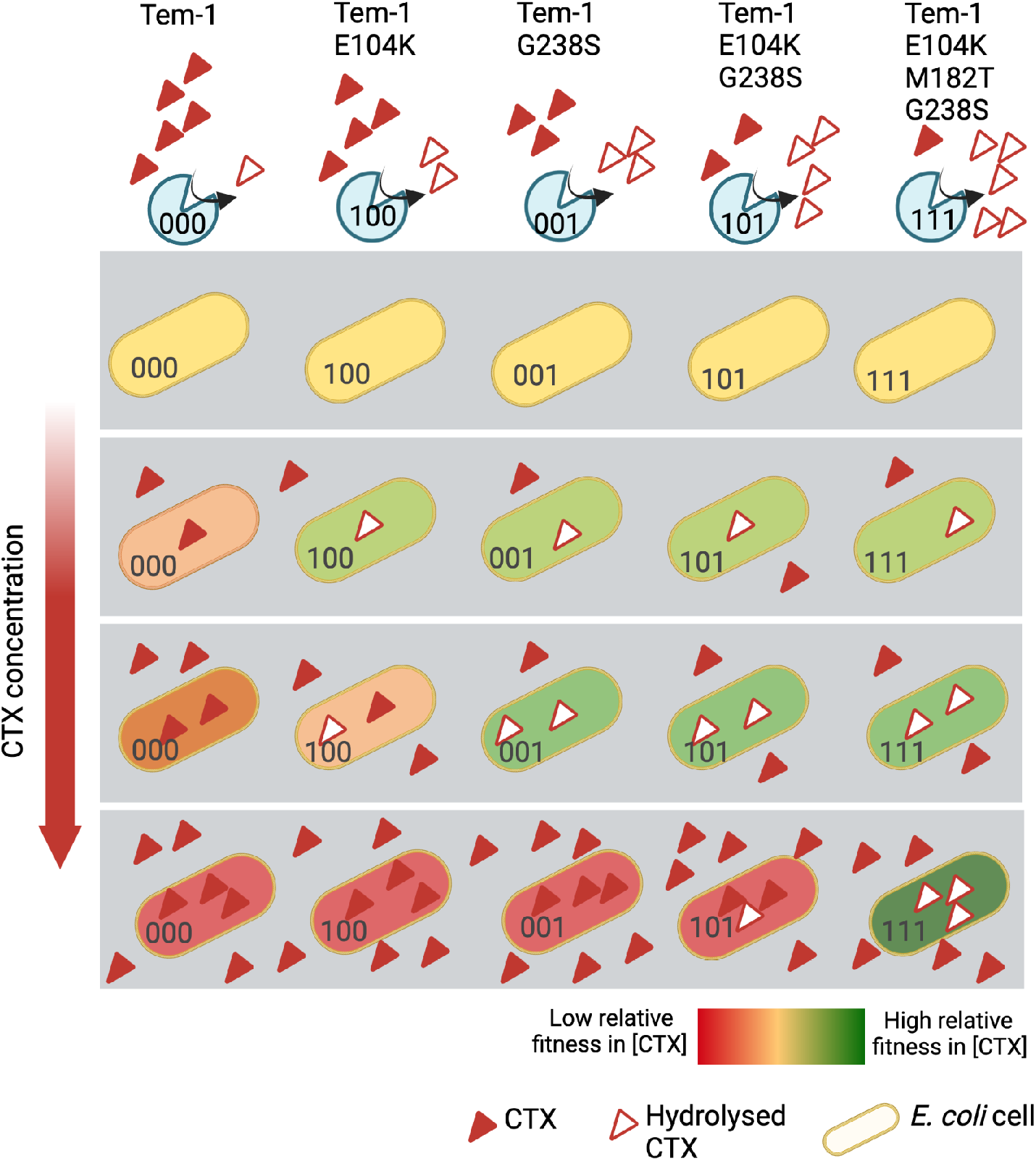
Proposed cause of equivalent fitness amongst competing genotypes at sub-inhibitory antibiotic concentrations. Presented is the relative fitness of five representative TEM-1 mutant genotypes competing with each other in particular concentrations of CTX (signified by a single gray box). At low and intermediate initial concentrations of CTX, more active TEM-1 mutants are able to hydrolyze CTX, effectively reducing the periplasmic concentrations to sub-inhibitory levels and allowing fitness to be maximal. This internal hydrolysis of CTX results in classes of types with equivalent fitness, while susceptible mutants with higher concentrations of CTX in the periplasm have lower fitness. As the concentration of CTX in the medium increases (see lower grey boxes), this class of fitness-equivalent mutants is predicted to decrease in number, leaving only those which can match the rate of diffusion of CTX into the cell with the rate of degradation. Note this figure presents the fate of cells at early stages of competition, before cells have a chance to alter the environmental concentrations of CTX. As the density of variants that can hydrolyze periplasmic CTX increases due to growth, eventually the environmental CTX will be degraded, removing the relative fitness difference of resistant and susceptible cells. If the density of high resistant variants increases rapidly and leads to a rapid degradation of environmental CTX, the time window in which differences in TEM-1 activity against CTX for low resistant variants (i.e., TEM-1 vs TEM-1 E104K) impact fitness may be small, leading to fitness stratification for the low resistance types.

Our observation of fitness stratification may help refine of our understanding of how concentrations of antibiotic below MIC – as found in environmental settings such as wastewater (28, 36-39) – may select for antibiotic resistant strains (13, 29, 40-43). While it is clear that antibiotic resistant genes confer a benefit in sub-inhibitory conditions (13, 29), little is known about the evolution of such genes in these conditions. This study provides grounds to doubt that sub-MIC environmental concentrations of antibiotics may select for mutations conferring the highest resistance. If many genotypes have equivalently high fitness above a threshold value of resistance, there are many factors that could determine which variants eventually predominate. For example, mutation bias (44), trade-offs between resistance and growth (45), and collateral sensitivity to other antibiotics (46, 47) could be important. Selection for resistant types in natural environments may be further complicated by the striking inverse relationship between resistance and fitness in spatially structured sub-MIC environments – a phenomenon which is associated with filamentation of susceptible types as previously reported (25). It is clear that a degree of spatial structuring is common to microbial populations in their natural habitats. It is thus interesting to consider whether environmental spatial structuring and selection at sub-inhibitory antibiotic concentrations may select for less resistant genotypes. The lack of selective benefit for genotypes such as TEM-52 in our study conflicts with the wide and frequent isolation of this mutant β-lactamase in clinical settings (48, 49). One scenario that is compatible with our observations is that (1) environmental conditions with sub-lethal antibiotic concentrations select for limited resistance, (2) upon migration of the ß-lactamase hosts to clinical environments – or other rare habitats with high antibiotic concentrations – there will be relatively short bouts of selection for high-resistant mutations, and (3) these variants are then maintained under the environmental conditions, where all high-resistance alleles have equivalent fitness and near neutral evolution predominates.

There are important considerations to make when applying these findings and ideas to real-world scenarios. For practical reasons, we used equal starting frequencies of the competitors, and we can only speculate how combining competitors at extreme frequencies – such as when mutations arise *denovo* – would affect resulting fitness differences. It is possible the environment-dependent fitness-benefits of genotypes may continue regardless of the competitors starting frequency, resulting in the eventual fixation of genotypes provided enough generations. Alternatively, there may be negative-frequency dependent interactions which establish an equilibria of high and low resistant types – possibly as a result of CTX degradation (21). It is also unclear whether the spatial-structure associated benefit of low-resistance genotypes is caused only by β-lactams such as CTX (this study) and carbenicillin (25), or whether this phenomenon extends to other antibiotics which cause milder changes in the cellular aspect-ratio of sensitive strains (50). Finally, although we have used strains with similar levels of fitness in the absence of antibiotics, we predict that increasing the cost of resistance may cause a negative correlation between resistance and fitness amongst high-resistance types (51). It would therefore be highly relevant to measure the relationship between fitness and resistance for fitness landscapes of either β-lactamase – or alternative resistance mechanisms – which involve higher fitness costs. We suggest that such experiments will help provide a fuller picture of how clinically relevant strains evolve and depend on specific selective conditions.

## Methods

### Strains, media and growth conditions

All strains used in this study derive from *Escherichia coli* MG1655 (see supplementary tables 1). *E coli* MG1655 *galk::SYFP2-cat* (DA28100) and *E coli* MG1655 *galk::mTagBFP2-cat* (DA28102) were kindly gifted by Peter A Lind (Uppsala University, Uppsala, Sweden). All experiments were performed in minimal media composed of the following: 8.5 g/L Na_2_HPO_4_.2H_2_O, 3 g/L KH_2_PO_4_, 0.5 g/L NaCl, 1 g/L NH4Cl, 4.93 g/L MgSO_4_.7H_2_O, 147 mg/L CaCl_2_.2H_2_O, 0.2 % (W/V) Casamino Acids (Difco), 2 mg/L Uracil, 1 µg/L Thiamine, 0.4% (W/V) Glucose, 2 mg/L Uracil, 1 µg/L Thiamine. Agar was added at 1.5 % (W/V) in order to make solid media and was autoclaved separately from other components. Expression of ß-Lactamase was induced by addition of 50uM isopropyl β-D-1-thiogalactopyranoside (IPTG) to the competition media. CTX solutions were prepared from Cefotaxime Sodium Salt (Sigma), diluted in minimal media solution and stored as stock solution at 5.12 mg/mL at −20 °C, before final dilution to stated working concentrations. Strains were prepared for flow cytometry by dilution in M9 salt solution (8.5 g/L Na_2_HPO_4_.2H_2_O, 3 g/L KH_2_PO_4_, 0.5 g/L NaCl, 1 g/L NH4Cl, 4.93 g/L MgSO_4_.7H_2_O) that was filtered using a 0.2 µm syringe filter. Strains were grown at 37 °C. Liquid cultures were always aerated with orbital shaking.

### Strain reconstruction

All TEM-1 alleles in this study were reconstructed into a clonal strain of *Escherichia coli* MG1655 featuring a fluorescent *bfp* reporter gene inserted into *galK* (galK::mTagBFP2-FRT) (29). TEM-1 alleles – derived from a previous study (5) – were amplified by PCR and inserted into a remaining section of *galK* using the ‘Quick and Easy *E. coli* Gene Deletion Kit’ (Gene Bridges), using ampicillin resistance conferred by the TEM-1 genes to select for double recombinants. Individual ampicillin resistant colonies were isolated, cryogenically stored, and amplicon sequenced to confirm the absence of additional mutations in TEM-1. Each biological replicate used in subsequent assays represents an individual transformant.

### MIC measurements

All MIC assays were performed in the identical minimal media used for competitions, using a similar inoculum as experimental competition in liquid media. Assays were initiated with overnight cultures of each biological replicate in minimal media, grown for approximately 18 h until stationary phase. Cultures were then subculture with a 1:100 dilution and grown for ∼3 hours until a final cell density of 1.5 × 10^8 /mL. Meanwhile 2mL deep-well 96-well plates were filled with 400uL of minimal media featuring IPTG and a 2-fold dilution series of CTX over 12 rows (25.6 to 0.0125 µg / mL of CTX, except for one row intended for the E104K M182T G238S genotype which ranged from 102.4 to 0.05 µg / mL of CTX). A further 1:10 dilution of the subculture was made was made and 3uL of each genotype was added to the wells of each column, resulting in final concentration of cells of 1.5 × 10^5 /mL in each well. Plates were sealed with Breathe-Easier Sealing Film (Diversified Biotech) and shaken with orbital shaking at 750rpm for 24 hours, whereupon 200uL from each well was transferred to transparent 96 well plates for OD600 measures by a Victor3 microtiter plate reader (PerkinElmer). Minimum inhibitory concentrations were interpreted as the lowest concentration of CTX maintained OD600 at less than 0.1. MIC values of individual genotypes are reported as median values, while MIC values of groups of genotypes are the means of the median MIC values.

### Pairwise fitness assays

Pairwise fitness assays involved direct competition of each stated genotype with a common *yfp*-expressing competitor (E coli MG1655 galK::SYFP2-FRT), performed in either liquid or on solid media. Liquid media competitions were initiated with overnight monocultures of each competitor grown in 2 mL of minimal media. Overnights were grown for ∼18 hours, then each competitor was subcultured together with a 1:200 dilution into fresh minimal media, and incubated for ∼ 3 hours. Meanwhile, the competition media was prepared – 794 µL of minimal media containing IPTG and specified concentrations of CTX (see results) were aliquoted into rows of a 2mL deep-well 96 well plate. Subcultures were then added to the competition media, by pipetting 6 µL of the subculture into to the competition media. To measure the initial ratio and concentration of cells, 50 µL was sampled from competition wells without CTX to avoid any possible affect of CTX on the ratio of each competitor). These samples were each added to 200 µL of filtered M9 salt solution inside the wells of a 96 well plate and BFP+ and YFP+ cellular events were quantified by flow cytometry using a Macsquant10 (Miltenyi biotec). The competition plates were then sealed with Breathe-Easier Sealing Film (Diversified Biotech), and incubated at 37C with 750rpm orbital shaking. After 24 and 48 hours of competition, 10 µL samples were taken from each well and diluted 1:500 in M9-salt solution to allow flow cytometry of the ratio and concentration of the competitors. Flow cytometry was performed using a medium flow rate using SSC 1.4 as a trigger (no secondary trigger) with 25,000 events counted per sample. All events measured as >1 by the 450/50 nm and 525/50 nm channels were respectively counted as BFP+ and YFP+ events. Fitness assays performed on solid media were performed similarly to the liquid-media competitions with the following distinctions. Overnight cultures of each competitor were subcultured together with a 1:1000 by dilution into fresh minimal media, incubated with shaking for 3 hours, and were then further diluted (1:10) and samples of this dilution were taken for cellular quantification by flow cytometry. A drop of 1 µL of the mixed culture was spotted onto minimal media solidified with agar (containing a relevant concentration of CTX). Spotting was performed twice to allow the destructive harvesting of colonies at 24 and 48 hours. Harvesting was performed by scooping each colony into the base of a sterile 1mL pipette tip, with the cells resuspended by vortexing the pipette tip in a microcentrifuge tube containing a 1mL M9 salt solution. The cell solution was then diluted 1:100 in M9 salt solution and the ratio and concentration of cells were quantified by flow cytometry. Selection coefficients were calculated as s = [ln(R(t)/R(0))]/[G] (where R is the ratio of a genotype relative to competitors at time (t) and G is the number of generations (Dykhuizen, 1990). The number of generations was calculated by ln(final population/initial population)/ln(2). Presented selection coefficient values for the genotypes were base-line adjusted to make the selective coefficient for TEM-1 equivalent to 0. All pairwise fitness assays were performed with three biological replicates each on a separate occasion.

### Four-genotype fitness assays

We measured the relative fitness of the four genotypes connected by a single mutation to each node of the TEM-1 to TEM-52 fitness landscape, as well as the relative fitness of all eight genotypes competed together. Competitions were performed in liquid or on solid media supplemented with either 0, 0.02 or 0.04 µg/mL CTX. For each replicated assay, all eight strains were cultured for ∼18h overnight in 2 mL minimal media. The strains were then mixed together by adding 2 µL of each culture in a 1:1:1:1 ratio into 2mL of minimal media (or into 4 mL in the case of the eight genotype competitions). Strains were then grown together for 3 hours and then diluted 1:10 in M9 salt solution and the total cellular number of BFP+ events were measured by flow cytometry as per the pairwise fitness assays (see above). Subcultures were then inoculated into the competitive media. Liquid media competitions were initiated with 2.4 uL of each mixed subculture inoculated into 2 mL of minimal media supplemented with IPTG and either 0, 0.02 or 0.04 µg/mL CTX. Solid media competitions were then initiated with a 1 µL drop of subculture spotted onto minimal media supplemented as per the liquid competitions. Competitions were founded with approximately 3 × 10^4^ cells mL^-1^ for liquid competitions and ∼ 3 × 10^4^ cells in the case of spatially structured competitions. The concentration of BFP+ cells in the subculture was further quantified by flow cytometry, and the mean value of these measures – before and after inoculation in the test media – was used as measures of the initial concentration of cells in each competition. A sample of 1mL of subculture was then added to a cryogenic tube supplemented with glycerol and flash frozen for subsequent amplicon sequencing (see below). Liquid and Solid media competitions were then incubated at 37 °C (with shaking for the liquid cultures) for 24 or 48 hrs until sampling. Sampling of Liquid cultures were performed by removing 150µL of competition media after 24 and 48 hrs. Glycerol was added to the sample which was then frozen at −70 °C for amplicon sequencing at a later date. An additional 10 µL sample of competition media was taken, diluted 1:400, and used to measure the concentration of BFP+ cells by flow cytometry (see Pairwise competition methods for flow cytometry details). Colonies that had grown in solid media after 24 and 48 hours were destructively harvested in the same way as the pairwise competitions (see methods above). Colonies were resuspended in 1mL of M9 salt solution and the samples stored and BFP+ events quantified in the same fashion as the Liquid media samples. A total of four biological replicates of each genotype was used in this experiment, with a different biological replicate competed on separate occasions and competition media freshly prepared for each replicate.

Simultaneous to these competitions, controls experiments were conducted on isogenic populations of each of the eight competitors. Instead of mixing each competitor with 3 other genotypes, isogenic subcultures were made that were otherwise treated identically to samples from four-genotype competitions. Flow cytometry measures were then taken to establish the degree of filamentation in each population subject to CTX. To do so, these control populations were measured after at 24 and 48 time periods using the FSC channel of a Macsquant10 (Miltenyi biotec). Flow cytometry was performed with the same setting used in pairwise fitness assays (see above), including gating of events >1 in the 450/50 nm channel to remove non-fluorescent particles. The arbitrary FSC value that defined the largest 1% of events in populations in CTX-free environments was established, and the percentage of samples that exceeded that FSC value was established for populations subject to CTX. Three biological replicates were conducted for each genotype and treatment.

### Amplicon sequencing and data analysis

Amplicon sequencing was used to measure the relative frequency of genotypes in the four and eight-genotype competitions to allow calculations of relative fitness. Primers were designed to produce a 475 bp amplicon which spanned the three mutation sites. Forward and reverse primers were each labelled with a 5’ four bp sequence to allow multiplexing. Each of the nine competitions (the eight four-genotype competitions and one eight-genotype competition) were amplified with a unique pair of indices for each treatment and each replicate (Table S3). Frozen samples from each time points were thawed and used directly as a template for PCR. Different volumes of sample were used as template from the 24 and 48 hr time points (1 µL) compared to intimal time points (2 µL). PCR reactions were performed on each occasion with the template of a different replicate. Reactions were performed in a total volume of approximately 30 µL, including 15 µL Phusion Flash High-Fidelity PCR Master Mix (Thermo Scientific), forward and reverse PCR primers to a final concentration of 0.5 µM, and the thawed template. PCR was performed over 30 cycles with an initial 5-minute incubation step to allow for cell lysis. Amplicons from each reaction were confirmed by electrophoresis. PCR products with differing indices from each treatment and replicate were then pooled together, purified with NucleoSpin Gel & PCR Clean-up (BIOKE). The TruSeq Library Preparation Kit (Illumina) was then used to prepare libraries for MiSeq PE-250 sequencing (Illumina), with a preparation done for each pool of nine samples. Library preparation and high throughput sequencing were performed by the Cologne Center for Genomics.

The sequencing data were analysed with CLC Genomics Workbench 11.0. We trimmed the sequences with “Trims Reads 2.1”, with standard settings except for quality limit = 0.001 (equivalent to Phred Score of 30) and minimum number of nucleotides = 260. All broken pairs were discarded at this step. Next, we mapped the trimmed reads to a set of reference sequences, representing all possible combinations of our custom barcodes and TEM alleles (i.e,, not only those combinations that were expected, but all possible permutations). For this step, “Map Reads to Reference” was used with standard settings, except length fraction = 0.995 and similarity fraction = 0.995 (i.e., the reads have to be perfect match to the reference to map). We then used the total read count mapped to each sequence as the estimate of that sequence’s frequency in the population. (Note that as an alternative approach, we first subdivided the data from each Illumina library based on our custom barcodes for different conditions and trimmed the barcodes, using Demultiplex Reads 1.0, and then trimmed and map these reads as above, but to reference sequences without the custom primers present. The resulting frequencies of alleles were nearly identical.)

We found unexpected combinations between TEM alleles and our custom barcodes at appreciable frequencies, with the three possible sequences with a single unexpected position in the TEM sequence being more common (mean frequency ±standard deviation = 0.023 ±0.019, determined over all nodes in the fitness landscape) than the sequence with two unexpected positions in the TEM sequence (0.003 ±0.002). These unexpected alleles therefore appear to have arisen due to recombination in the first PCR step. We know which real alleles were in the inoculum, and each allele other than the “nodal progenitor” (i.e., TEM-1 for the competition between TEM-1 and all three one-step mutants) has a unique mutation that allows us to determine its frequency from the number of (essentially full length) sequencing reads that mapped to each of the eight TEM alleles. The frequency of the nodal progenitor is then one minus the sum of the three other genotypes. If the estimated frequency of the nodal progenitor was very low (< 0.001), we simply used the relative frequency of reads that mapped to the nodal type as an approximation of its frequency. Genotype frequencies were then converted into selection coefficient values using the same method described in ‘pairwise fitness assays’ (see above).

### Models of fitness

We explored three simple models predicting resistance from differences in MIC or from the presence of mutations in TEM. Models 1a and 1b are based on the difference in log[MIC] for a genotype compared to the mean of the four genotypes corresponding to a node (*ΔMIC*). Model 1a assumes a linear relationship between ΔMIC and the predicted selection coefficient *S*, such that for the *i*^th^ allele *S*_*i*_ = *α* · *ΔMIC*_*i*_, where *α* is a constant estimated from the data for each experimental condition. Model 1b assumes a linear relationship that is constrained to a minimum *γ*_min_, such that if *α* · *ΔMIC*_"_ > *γ*_min._ Then *S*_*i*_ = *γ*_min_, as well as a maximum *γ*_max_. Model 1a has one free parameter (*α*), whereas Model 1b has three parameters (*α, γ*_min_ and *γ*_max_). Model 2 assigns alleles to two classes depending on the presence of the mutations E104K, M182T or G238S, and assigns each fixed selection coefficients *S*_*1*_ and *S*_*2*_. Model parameters were estimated by minimizing the negative log likelihood (NLL) with grid searches over a broad parameter space, and the NLL was calculated from the residual sum of squares (52).

### Simulations of serial passaging

To evaluate multi-passage evolution on this landscape, for each condition we performed 1000 independent 100-passage simulations, with the ancestral TEM as the starting genotype. Each passage is initiated by the initial genotype or that sampled in the previous passage, together with the three genotypes with a Hamming distance = 1 in the landscape. The final frequency *f* of a genotype *i* is *f*_*i*_ = (1 + *S*_*i*_)^*g*^, where *S*_*i*_ is the empirically estimated selection coefficient for that genotype and *g* is the number of generations for each passage. Based on the cell counts from flow cytometry, we estimated 15.77 generations had occurred by 48 h, a number that was consistent over the different antibiotic and media treatments (standard error of the mean = 0.21). We therefore assumed *g* = 16 for all simulations. At the end of growth, one individual is randomly chosen using a pseudorandom number generator (*sample*() in R) to initiate the next passage, with the probability of sampling being weighed by the normalized frequency of each genotype. After 100 passages, we consider the final genotype selected as the evolutionary endpoint for that replicate.

### Code and data accessibility

Raw sequence reads for the four-genotype competitions are available via NCBI bioproject accession PRJNA844400. Scripts used for the presentation of fitness data (Fig. 3 and 6), for the analysis of flow cytometry data (Fig. 5A), for statistical models of fitness (Table 2) and for simulations (Fig. 6) are available via DOI:10.5281/zenodo.6818118. Supplementary data and calculations of MIC measures (Fig. 1 and Table 1), selection coefficients of pair-wise competitions (Fig. 2), selection coefficients of four-genotype competitions (Fig. 3 and 4) and filamentation measures (Fig. 5) are similarly available at via DOI:10.5281/zenodo.6818118.

## Acknowledgements

We would like to thank Dan Andersson, Erik Wistrand-Yuen and Peter A. Lind for the gift of the ancestral MG1655 strains and for related advice, and Marjon de Vos, Naama Brenner and Lukas Geyrhofer for valuable discussions, Loukas Theodosiou for statistical advice. Fig. 1C and Fig. 7 was created with BioRender.com. This study was made possible by the Human Frontiers Science Program grant RGP0010/2015.

## Supplementary Figures and Tables

**Supplementary figure S1:**
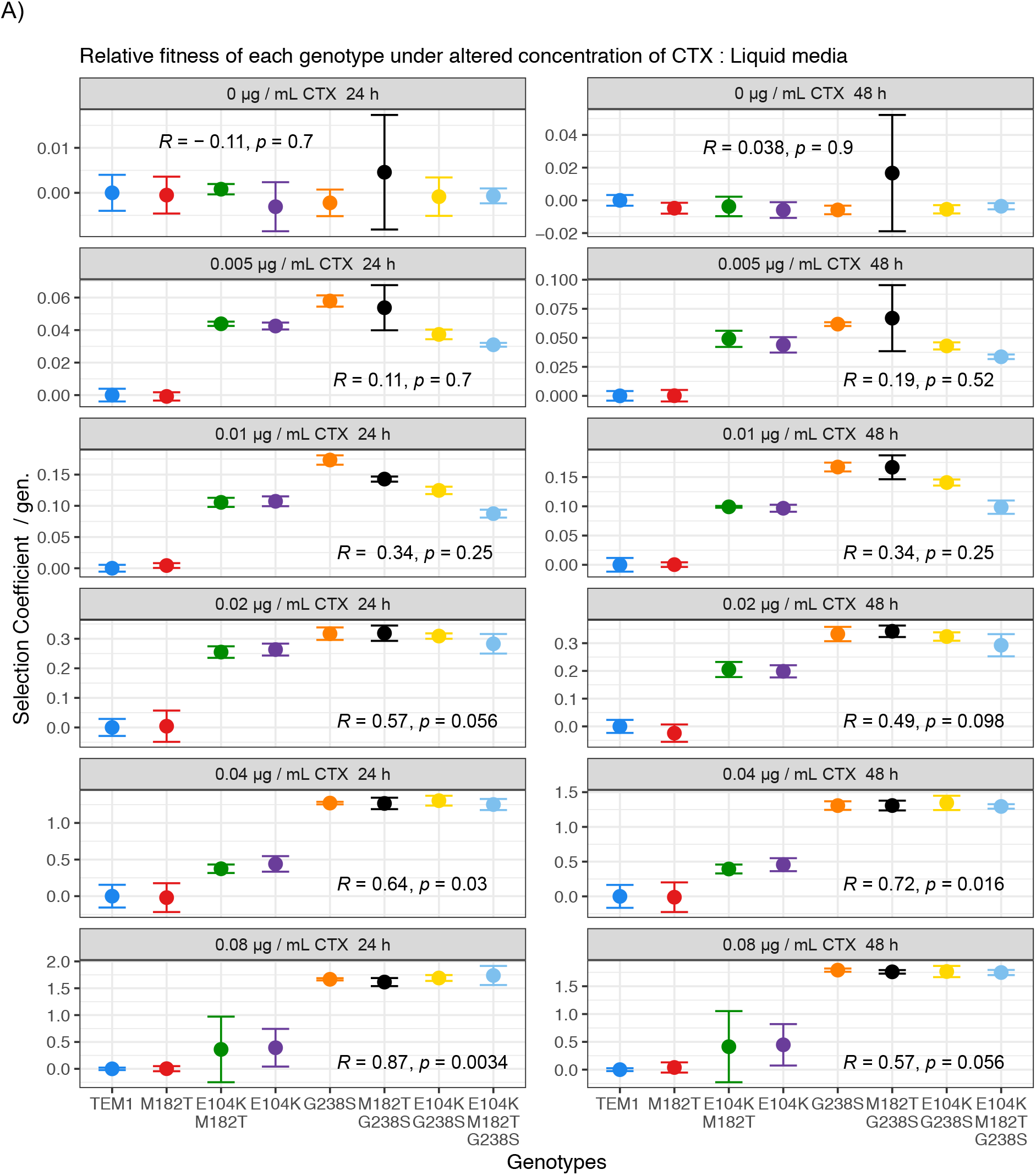
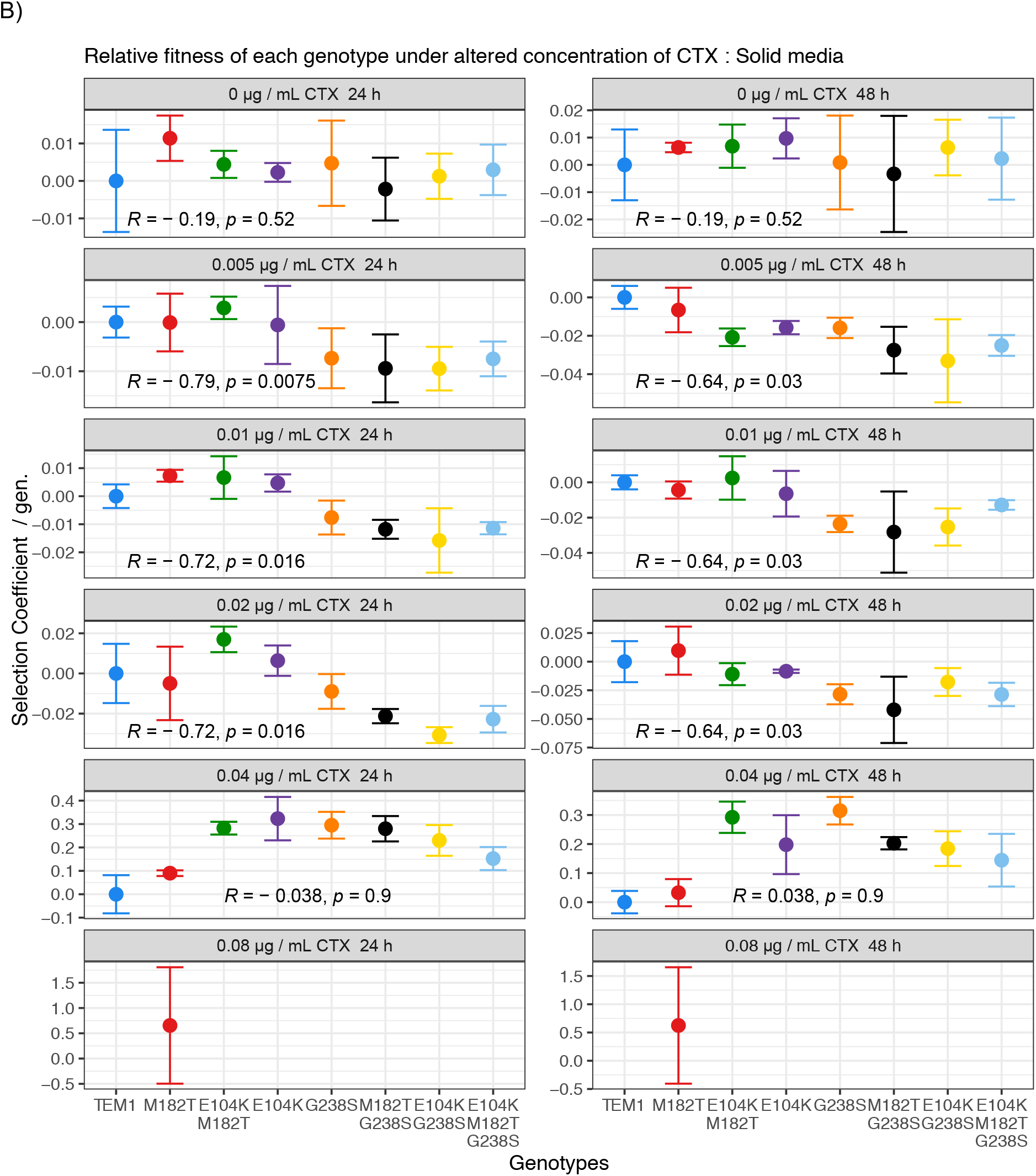
Pair-wise fitness assays relative to MG1655-YFP of the 8 BFP-labelled genotypes subject to varying concentrations of CTX. Competitions on solid media with 0.08 µg/mL of CTX resulted in null cell counts of either competitor for at least one replicate, and resulting means were not plotted. Data points represent the means of three biological replicates, error bars represent standard deviations and statistical tests presented are Kendall rank correlation coefficient Rho and p-values.

**Supplementary Figure S2:**
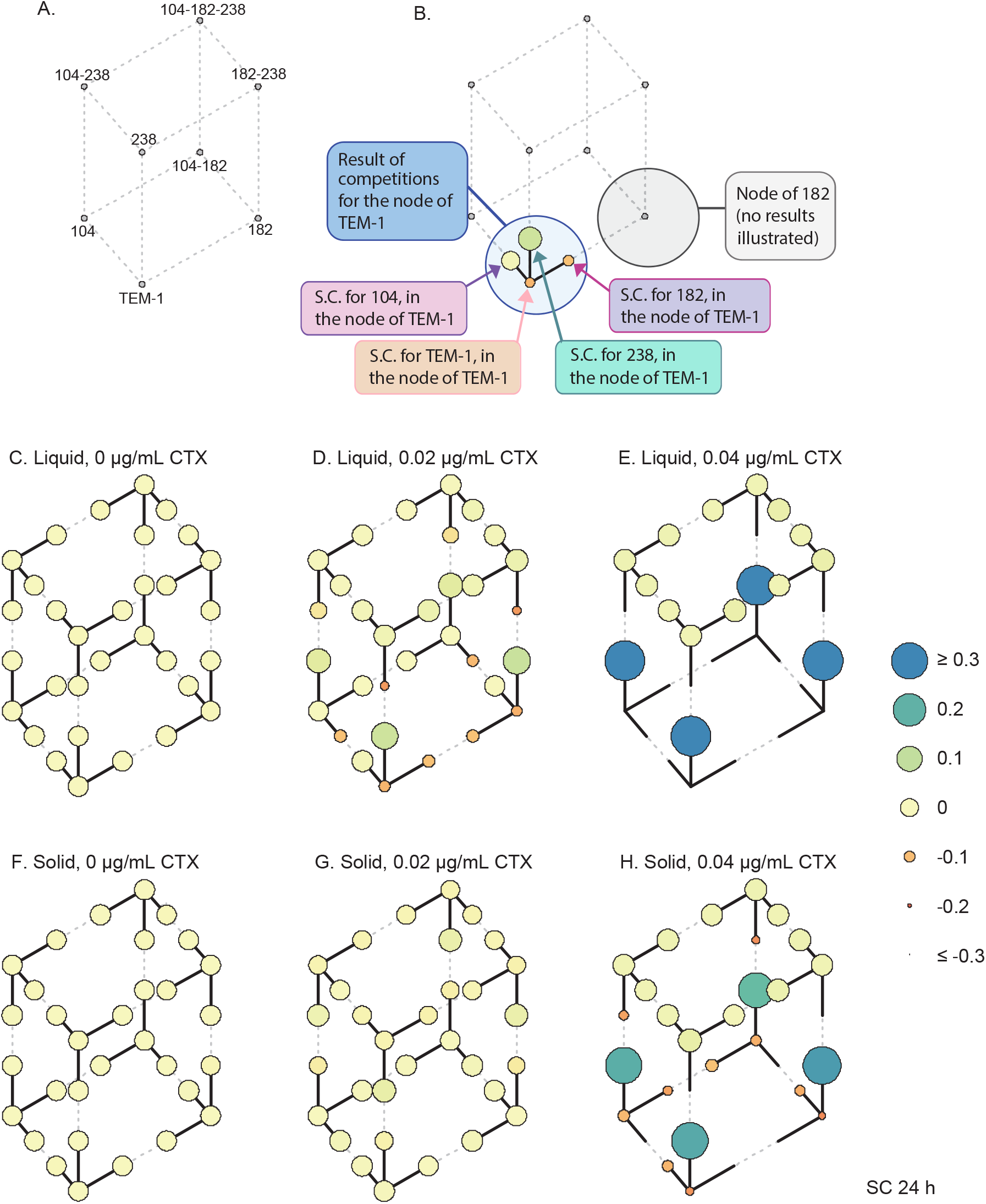
Mean selection coefficient (SC) measures of each genotype relative to three competitors at 24 hours. Figure is identical to Figure 3, however this supplementary figure presents the earlier time point of 24 hours. Each arrow or circle represents the SC of four biological replicates. Measures of error are not included to aid visual clarity.

**Supplementary figure S3:**
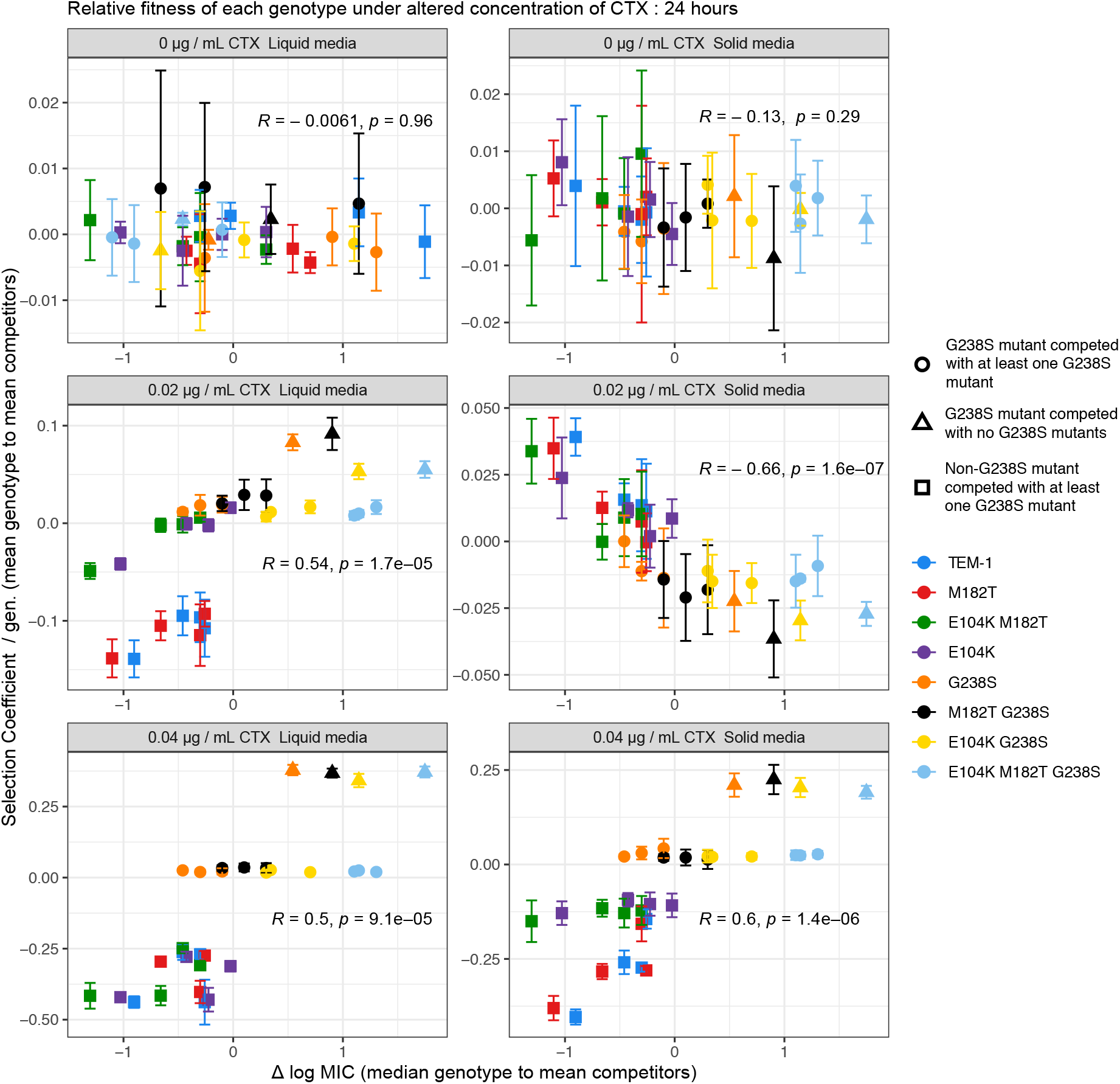
Concentration dependent relationships of resistance and selection coefficient of genotypes competed in 4-genotype competitions over 24 hours. The relationship at 24 hours is smiliar to that observed at 48 hours, except at 24 hours there is no significant correlation between selection coefficient and resistance in the absence of CTX. Data points represent the means of four biological replicates, error bars represent standard deviations and statistical tests presented are Kendall rank correlation coefficient Rho and p-values.

**Supplementary figure S4:**
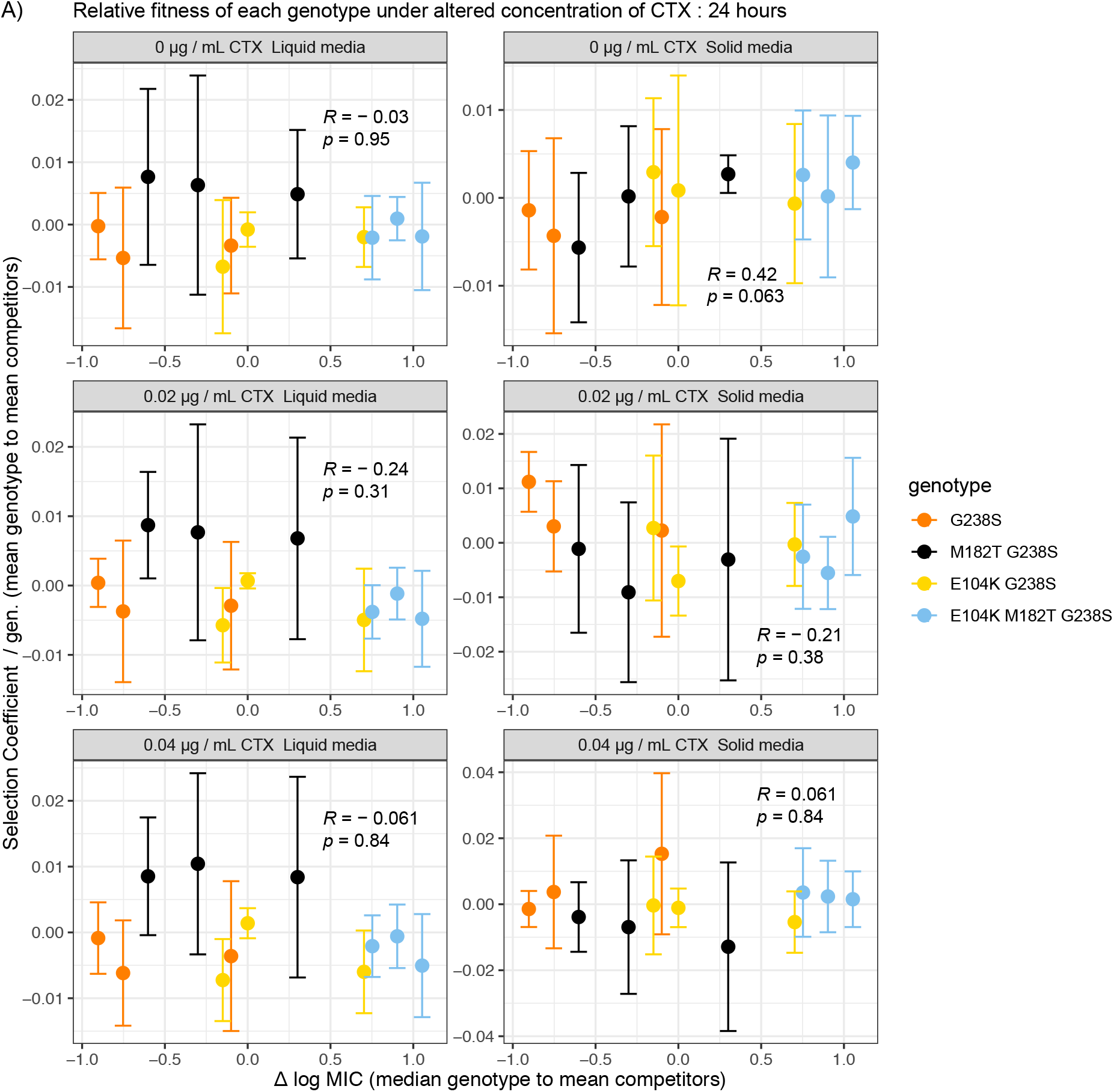
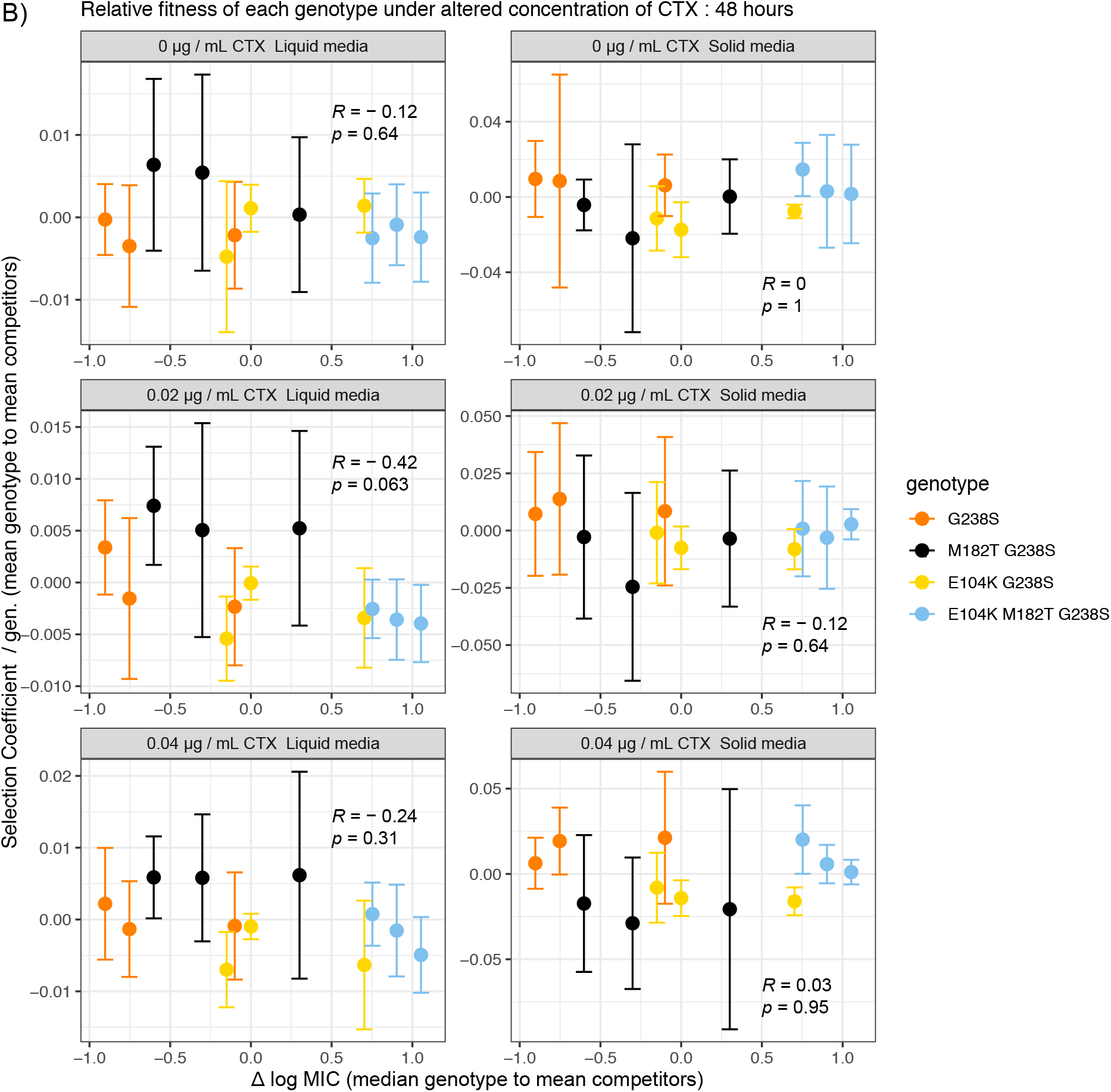
The relationship of resistance and selection coefficient of genotypes expressing a G238S mutation competed in 4-genotype competitions over A) 24 and B) 48 hours. Kendal rank correlations indicate no significant relationships between Selection coefficient and resistance under any condition for these genotypes. Data points represent the means of four biological replicates, error bars represent standard deviations and statistical tests presented are Kendall rank correlation coefficient Rho and p-values.

**Table S1:**
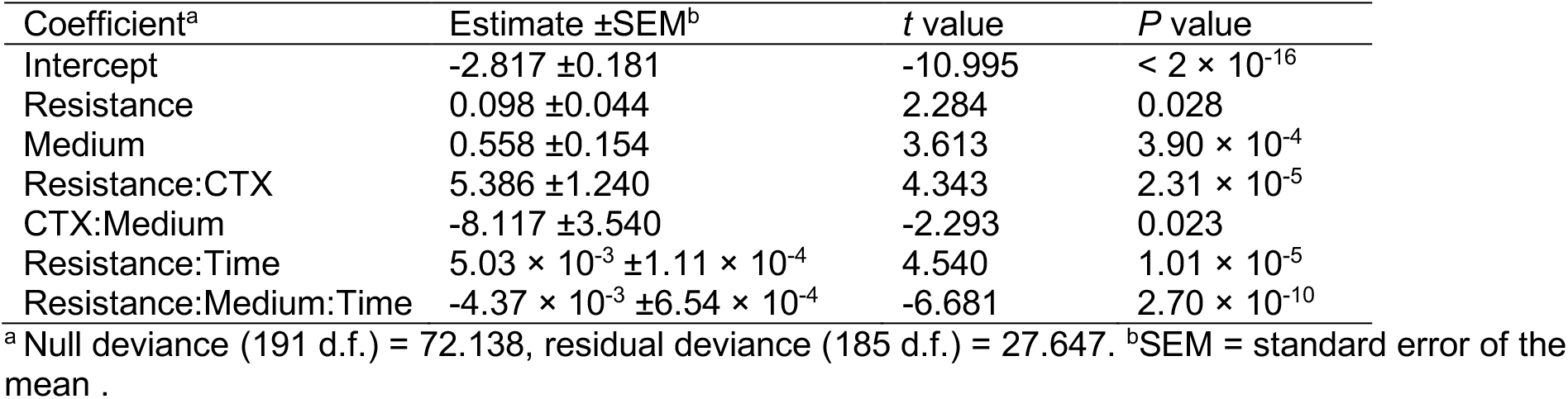
General linear model (GLM) for flow cytometry data. We analysed the log_10_-transformed fraction of FSC-A events which were higher than the 99 percentile for control populations with no antibiotics with a GLM. Variables included in the model were the ranked resistance of the genotype based on MIC (Resistance), liquid or solid medium (Medium), whether the population was exposed to low (0.02 μg/mL) and high (0.04 μg/mL) antibiotic concentrations (CTX), and the time (24 and 48) post inoculation (Time). The full factorial model gave an Akaike information criterion (AIC) value of 196.06, whereas a model with only two-way interactions resulted in AIC of 201.62. To further improve the model with two-way interactions, we dropped the two insignificant factors from this model (Time and the CTX:Time interaction), resulting in a slight improvement in model support (AIC = 199.28). Both the Resistance:Medium and Medium:Time interactions were highly significant, and so we combined these into a single three-way interaction, resulting in a final model with appreciably better support then the full factorial model (AIC 188.78). The final model is therefore Resistance + Medium + Resistance:CTX + CTX:Medium + Resistance:Time + Resistance:Medium:Time.

| Coefficient<sup>a</sup> | Estimate ±SEM<sup>b</sup> | *t* value | *P* value |
| --- | --- | --- | --- |
| Intercept | -2.817 ±0.181 | -10.995 | < 2 × 10<sup>-16</sup> |
| Resistance | 0.098 ±0.044 | 2.284 | 0.028 |
| Medium | 0.558 ±0.154 | 3.613 | 3.90 × 10<sup>-4</sup> |
| Resistance:CTX | 5.386 ±1.240 | 4.343 | 2.31 × 10<sup>-5</sup> |
| CTX:Medium | -8.117 ±3.540 | -2.293 | 0.023 |
| Resistance:Time | 5.03 × 10<sup>-3</sup> ±1.11 × 10<sup>-4</sup> | 4.540 | 1.01 × 10<sup>-5</sup> |
| Resistance:Medium:Time | -4.37 × 10<sup>-3</sup> ±6.54 × 10<sup>-4</sup> | -6.681 | 2.70 × 10<sup>-10</sup> |

<sup>a</sup> Null deviance (191 d.f.) = 72.138, residual deviance (185 d.f.) = 27.647. <sup>b</sup>SEM = standard error of the mean.

**Table S2:**
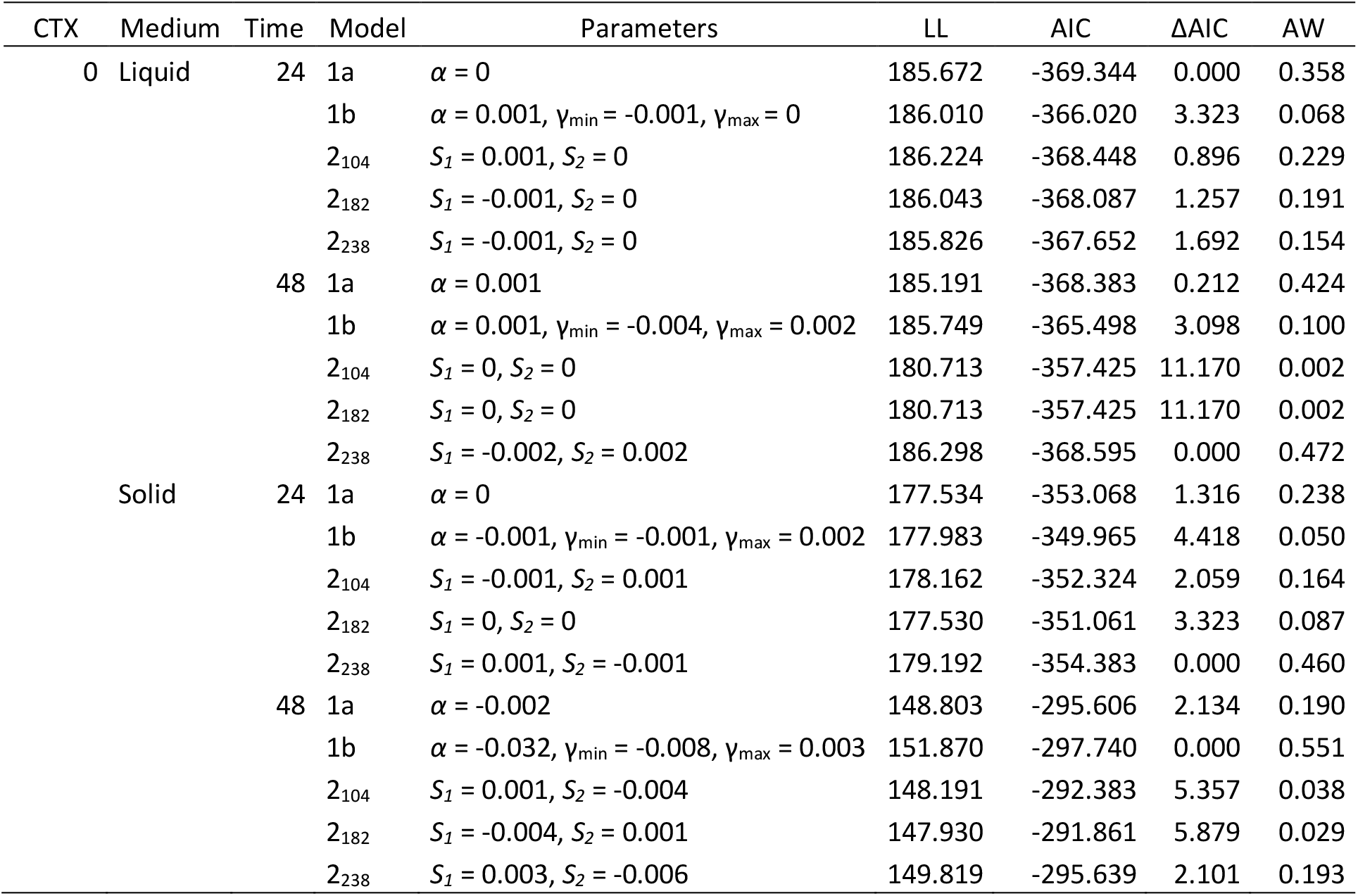
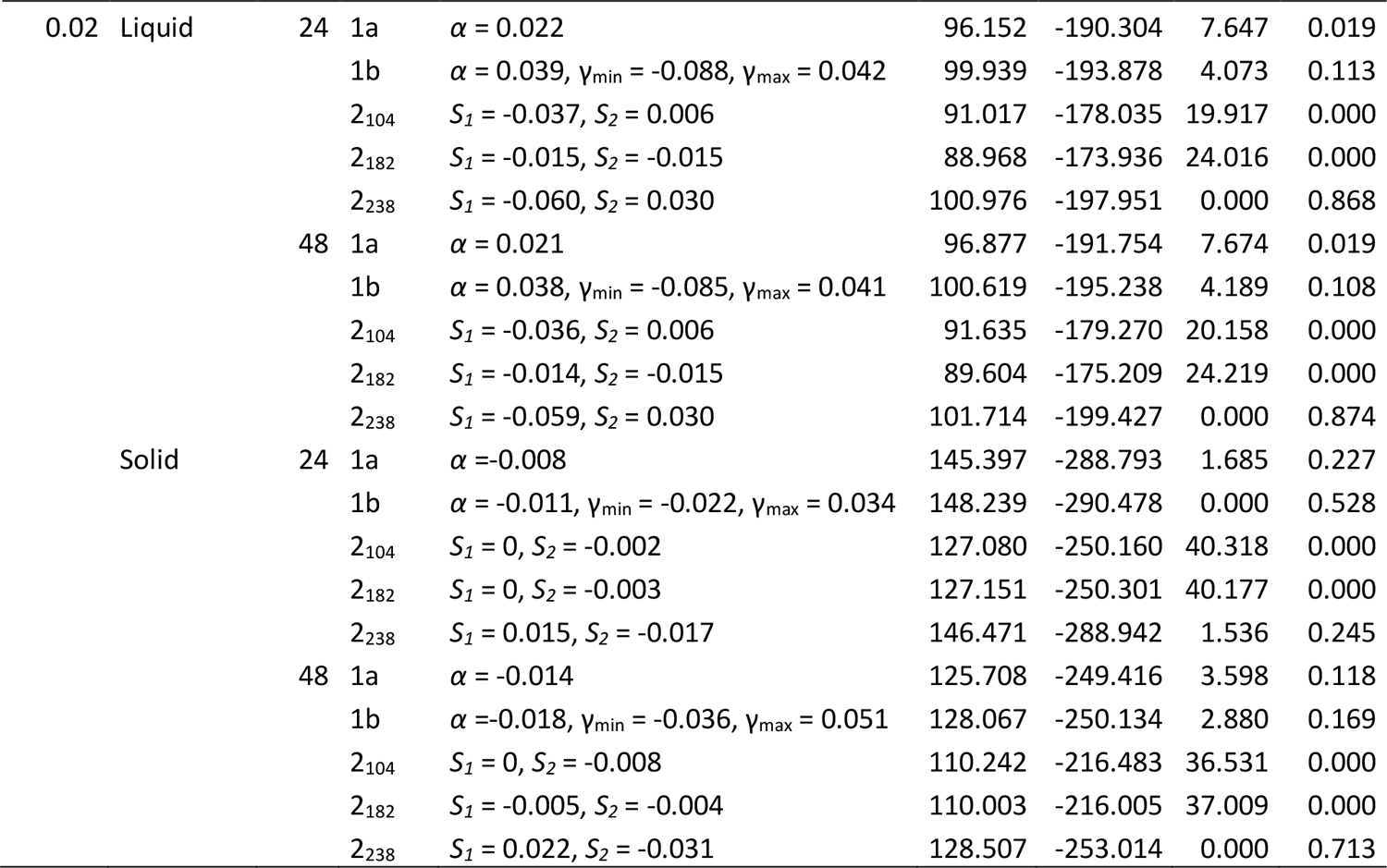
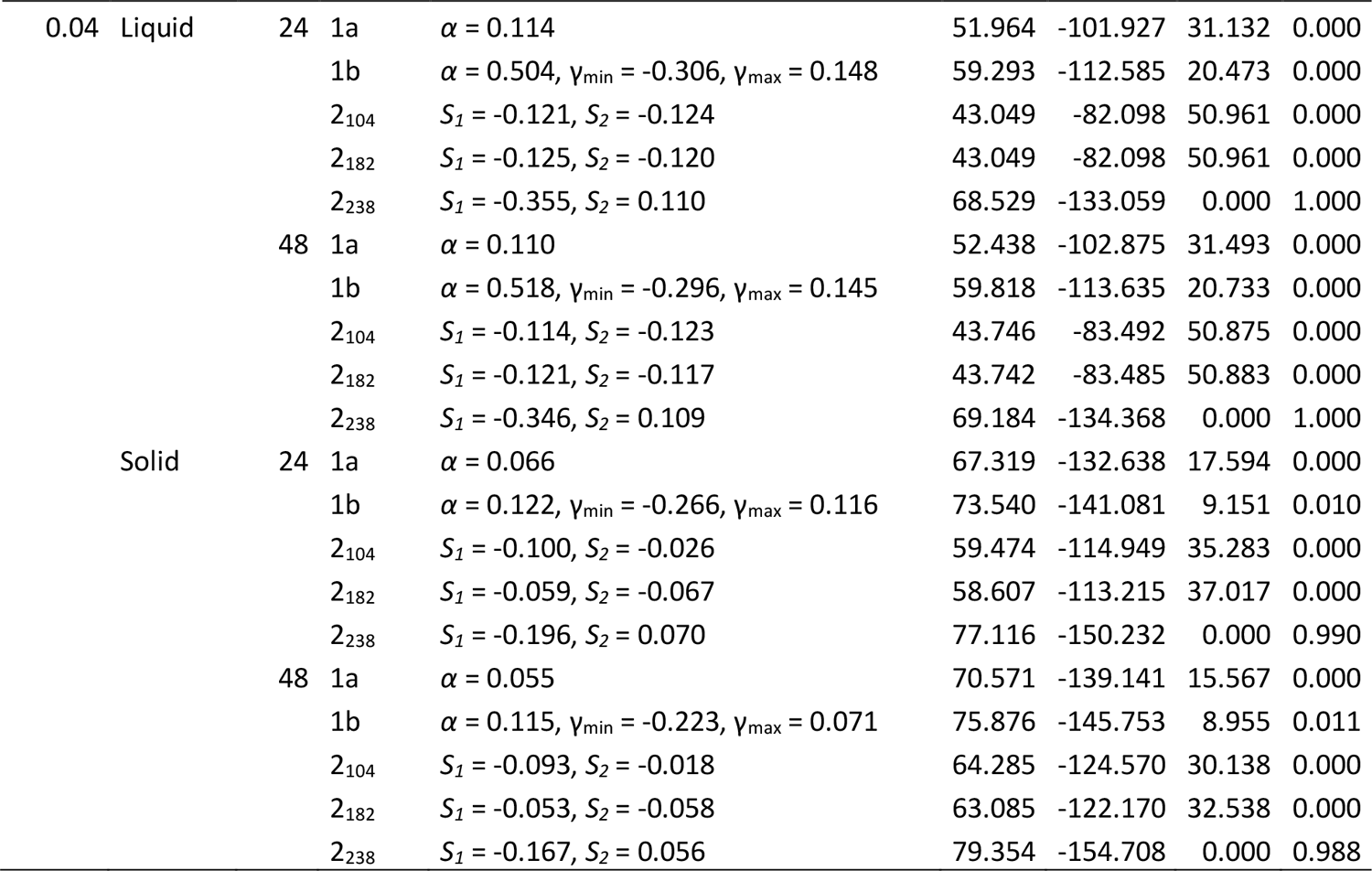
Model selection results. For each experimental condition (CTX concentration in μg/mL, medium type and time of sampling), the five models were fitted. For model 2, the subscripts indicate the site within the TEM gene used to classify the genotypes. Estimated model parameters are given (see Methods section for an explanation of model parameters), as well as the log likelihood, Akaike information criterion (AIC), the difference in AIC compared to the best supported model (ΔAIC), and the likelihood that a model is the best supported model within the set of models tested (Akaike Weight, AW). The table is divided over three pages, with results for experimental conditions with antibiotics on the next pages. Note that because we obtained a small residual sum of squares, log likelihood values are positive and AIC values are negative. Overall, Model 2_238_ is the best-supported model over the different datasets. Note that for intermediate CTX concentrations (0.02 μg/mL) on solid media, estimated model parameters indicate that high resistance alleles appear to be disadvantaged.

| CTX | Medium | Time | Model | Parameters | LL | AIC | ΔAIC | AW |
| --- | --- | --- | --- | --- | --- | --- | --- | --- |
| 0 | Liquid | 24 | 1a | $\alpha = 0$ | 185.672 | -369.344 | 0.000 | 0.358 |
| | | | 1b | $\alpha = 0.001, \gamma_{\min} = -0.001, \gamma_{\max} = 0$ | 186.010 | -366.020 | 3.323 | 0.068 |
| | | | 2<sub>104</sub> | $S_1 = 0.001, S_2 = 0$ | 186.224 | -368.448 | 0.896 | 0.229 |
| | | | 2<sub>182</sub> | $S_1 = -0.001, S_2 = 0$ | 186.043 | -368.087 | 1.257 | 0.191 |
| | | | 2<sub>238</sub> | $S_1 = -0.001, S_2 = 0$ | 185.826 | -367.652 | 1.692 | 0.154 |
| | | 48 | 1a | $\alpha = 0.001$ | 185.191 | -368.383 | 0.212 | 0.424 |
| | | | 1b | $\alpha = 0.001, \gamma_{\min} = -0.004, \gamma_{\max} = 0.002$ | 185.749 | -365.498 | 3.098 | 0.100 |
| | | | 2<sub>104</sub> | $S_1 = 0, S_2 = 0$ | 180.713 | -357.425 | 11.170 | 0.002 |
| | | | 2<sub>182</sub> | $S_1 = 0, S_2 = 0$ | 180.713 | -357.425 | 11.170 | 0.002 |
| | | | 2<sub>238</sub> | $S_1 = -0.002, S_2 = 0.002$ | 186.298 | -368.595 | 0.000 | 0.472 |
| | Solid | 24 | 1a | $\alpha = 0$ | 177.534 | -353.068 | 1.316 | 0.238 |
| | | | 1b | $\alpha = -0.001, \gamma_{\min} = -0.001, \gamma_{\max} = 0.002$ | 177.983 | -349.965 | 4.418 | 0.050 |
| | | | 2<sub>104</sub> | $S_1 = -0.001, S_2 = 0.001$ | 178.162 | -352.324 | 2.059 | 0.164 |
| | | | 2<sub>182</sub> | $S_1 = 0, S_2 = 0$ | 177.530 | -351.061 | 3.323 | 0.087 |
| | | | 2<sub>238</sub> | $S_1 = 0.001, S_2 = -0.001$ | 179.192 | -354.383 | 0.000 | 0.460 |
| | | 48 | 1a | $\alpha = -0.002$ | 148.803 | -295.606 | 2.134 | 0.190 |
| | | | 1b | $\alpha = -0.032, \gamma_{\min} = -0.008, \gamma_{\max} = 0.003$ | 151.870 | -297.740 | 0.000 | 0.551 |
| | | | 2<sub>104</sub> | $S_1 = 0.001, S_2 = -0.004$ | 148.191 | -292.383 | 5.357 | 0.038 |
| | | | 2<sub>182</sub> | $S_1 = -0.004, S_2 = 0.001$ | 147.930 | -291.861 | 5.879 | 0.029 |
| | | | 2<sub>238</sub> | $S_1 = 0.003, S_2 = -0.006$ | 149.819 | -295.639 | 2.101 | 0.193 |

**Table S2: Model selection results (continued section 2).**
| CTX | Medium | Time | Model | Parameters | LL | AIC | ΔAIC | AW |
| --- | --- | --- | --- | --- | --- | --- | --- | --- |
| 0.02 | Liquid | 24 | 1a | $\alpha = 0.022$ | 96.152 | -190.304 | 7.647 | 0.019 |
| | | | 1b | $\alpha = 0.039, \gamma_{\min} = -0.088, \gamma_{\max} = 0.042$ | 99.939 | -193.878 | 4.073 | 0.113 |
| | | | 2<sub>104</sub> | $S_1 = -0.037, S_2 = 0.006$ | 91.017 | -178.035 | 19.917 | 0.000 |
| | | | 2<sub>182</sub> | $S_1 = -0.015, S_2 = -0.015$ | 88.968 | -173.936 | 24.016 | 0.000 |
| | | | 2<sub>238</sub> | $S_1 = -0.060, S_2 = 0.030$ | 100.976 | -197.951 | 0.000 | 0.868 |
| | | 48 | 1a | $\alpha = 0.021$ | 96.877 | -191.754 | 7.674 | 0.019 |
| | | | 1b | $\alpha = 0.038, \gamma_{\min} = -0.085, \gamma_{\max} = 0.041$ | 100.619 | -195.238 | 4.189 | 0.108 |
| | | | 2<sub>104</sub> | $S_1 = -0.036, S_2 = 0.006$ | 91.635 | -179.270 | 20.158 | 0.000 |
| | | | 2<sub>182</sub> | $S_1 = -0.014, S_2 = -0.015$ | 89.604 | -175.209 | 24.219 | 0.000 |
| | | | 2<sub>238</sub> | $S_1 = -0.059, S_2 = 0.030$ | 101.714 | -199.427 | 0.000 | 0.874 |
| | Solid | 24 | 1a | $\alpha = -0.008$ | 145.397 | -288.793 | 1.685 | 0.227 |
| | | | 1b | $\alpha = -0.011, \gamma_{\min} = -0.022, \gamma_{\max} = 0.034$ | 148.239 | -290.478 | 0.000 | 0.528 |
| | | | 2<sub>104</sub> | $S_1 = 0, S_2 = -0.002$ | 127.080 | -250.160 | 40.318 | 0.000 |
| | | | 2<sub>182</sub> | $S_1 = 0, S_2 = -0.003$ | 127.151 | -250.301 | 40.177 | 0.000 |
| | | | 2<sub>238</sub> | $S_1 = 0.015, S_2 = -0.017$ | 146.471 | -288.942 | 1.536 | 0.245 |
| | | 48 | 1a | $\alpha = -0.014$ | 125.708 | -249.416 | 3.598 | 0.118 |
| | | | 1b | $\alpha = -0.018, \gamma_{\min} = -0.036, \gamma_{\max} = 0.051$ | 128.067 | -250.134 | 2.880 | 0.169 |
| | | | 2<sub>104</sub> | $S_1 = 0, S_2 = -0.008$ | 110.242 | -216.483 | 36.531 | 0.000 |
| | | | 2<sub>182</sub> | $S_1 = -0.005, S_2 = -0.004$ | 110.003 | -216.005 | 37.009 | 0.000 |
| | | | 2<sub>238</sub> | $S_1 = 0.022, S_2 = -0.031$ | 128.507 | -253.014 | 0.000 | 0.713 |

**Table S2: Model selection results (continued section 3).**
| CTX | Medium | Time | Model | Parameters | LL | AIC | ΔAIC | AW |
| --- | --- | --- | --- | --- | --- | --- | --- | --- |
| 0.04 | Liquid | 24 | 1a | $\alpha = 0.114$ | 51.964 | -101.927 | 31.132 | 0.000 |
| | | | 1b | $\alpha = 0.504, \gamma_{\min} = -0.306, \gamma_{\max} = 0.148$ | 59.293 | -112.585 | 20.473 | 0.000 |
| | | | 2<sub>104</sub> | $S_1 = -0.121, S_2 = -0.124$ | 43.049 | -82.098 | 50.961 | 0.000 |
| | | | 2<sub>182</sub> | $S_1 = -0.125, S_2 = -0.120$ | 43.049 | -82.098 | 50.961 | 0.000 |
| | | | 2<sub>238</sub> | $S_1 = -0.355, S_2 = 0.110$ | 68.529 | -133.059 | 0.000 | 1.000 |
| | | 48 | 1a | $\alpha = 0.110$ | 52.438 | -102.875 | 31.493 | 0.000 |
| | | | 1b | $\alpha = 0.518, \gamma_{\min} = -0.296, \gamma_{\max} = 0.145$ | 59.818 | -113.635 | 20.733 | 0.000 |
| | | | 2<sub>104</sub> | $S_1 = -0.114, S_2 = -0.123$ | 43.746 | -83.492 | 50.875 | 0.000 |
| | | | 2<sub>182</sub> | $S_1 = -0.121, S_2 = -0.117$ | 43.742 | -83.485 | 50.883 | 0.000 |
| | | | 2<sub>238</sub> | $S_1 = -0.346, S_2 = 0.109$ | 69.184 | -134.368 | 0.000 | 1.000 |
| | Solid | 24 | 1a | $\alpha = 0.066$ | 67.319 | -132.638 | 17.594 | 0.000 |
| | | | 1b | $\alpha = 0.122, \gamma_{\min} = -0.266, \gamma_{\max} = 0.116$ | 73.540 | -141.081 | 9.151 | 0.010 |
| | | | 2<sub>104</sub> | $S_1 = -0.100, S_2 = -0.026$ | 59.474 | -114.949 | 35.283 | 0.000 |
| | | | 2<sub>182</sub> | $S_1 = -0.059, S_2 = -0.067$ | 58.607 | -113.215 | 37.017 | 0.000 |
| | | | 2<sub>238</sub> | $S_1 = -0.196, S_2 = 0.070$ | 77.116 | -150.232 | 0.000 | 0.990 |
| | | 48 | 1a | $\alpha = 0.055$ | 70.571 | -139.141 | 15.567 | 0.000 |
| | | | 1b | $\alpha = 0.115, \gamma_{\min} = -0.223, \gamma_{\max} = 0.071$ | 75.876 | -145.753 | 8.955 | 0.011 |
| | | | 2<sub>104</sub> | $S_1 = -0.093, S_2 = -0.018$ | 64.285 | -124.570 | 30.138 | 0.000 |
| | | | 2<sub>182</sub> | $S_1 = -0.053, S_2 = -0.058$ | 63.085 | -122.170 | 32.538 | 0.000 |
| | | | 2<sub>238</sub> | $S_1 = -0.167, S_2 = 0.056$ | 79.354 | -154.708 | 0.000 | 0.988 |

**Table S3:**
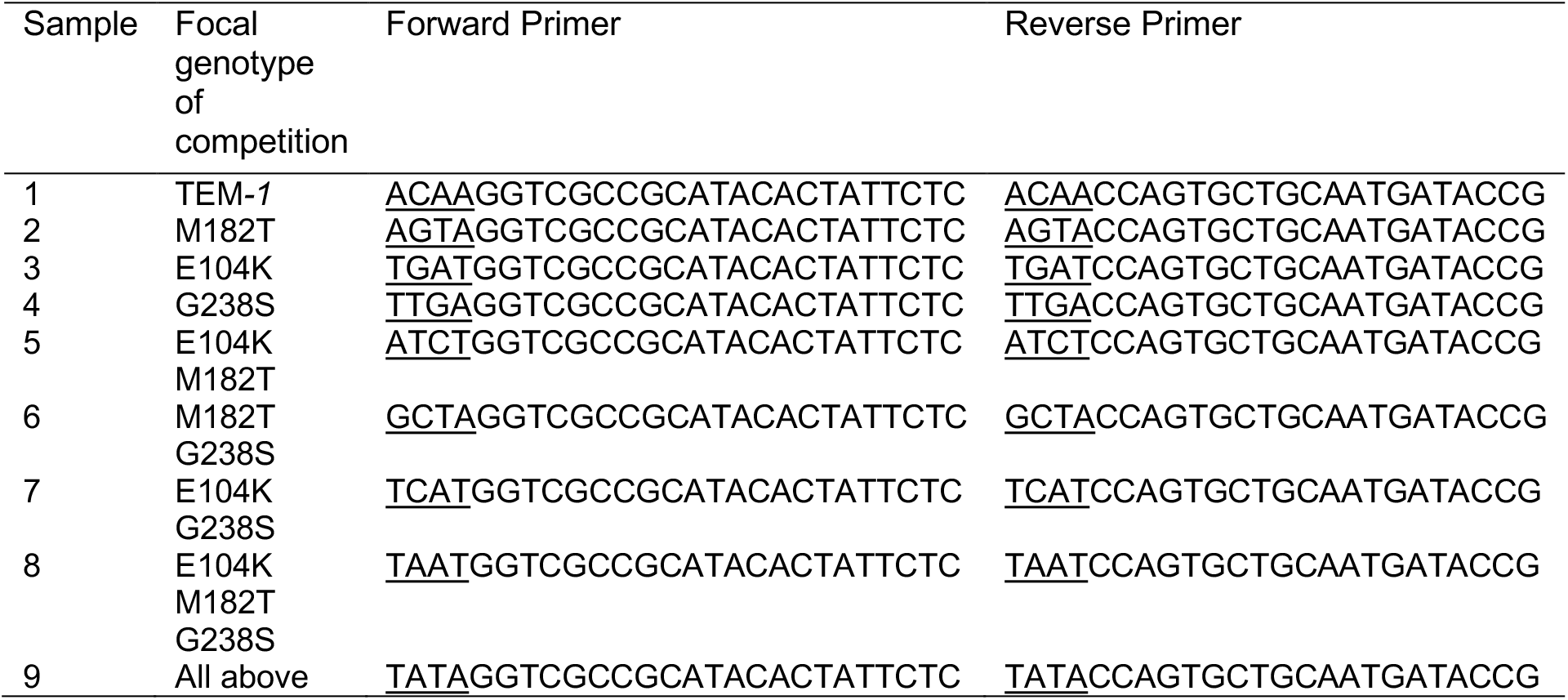
Primers used during amplicon sequencing. The underlined sequence (the first four 5’ nucleotides) is the barcode used to allow demultiplexing and allocation of reads to the original sample. Amplicons made with the below primers were pooled prior to Truseq library preparation.

## References

1. Wright S. 1932. The roles of mutation, inbreeding, crossbreeding, and selection in evolution. Proceedings of the XI International Congress of Genetics 1:356–66.

2. Lozovsky ER, Chookajorn T, Brown KM, Imwong M, Shaw PJ, Kamchonwongpaisan S, Neafsey DE, Weinreich DM, Hartl DL. 2009. Stepwise acquisition of pyrimethamine resistance in the malaria parasite. Proc Natl Acad Sci U S A 106:12025–30.

3. Weinreich DM, Delaney NF, Depristo MA, Hartl DL. 2006. Darwinian evolution can follow only very few mutational paths to fitter proteins. Science 312:111–4.

4. Hietpas RT, Jensen JD, Bolon DN. 2011. Experimental illumination of a fitness landscape. Proc Natl Acad Sci U S A 108:7896–901.

5. Schenk MF, Szendro IG, Salverda ML, Krug J, de Visser JA. 2013. Patterns of epistasis between beneficial mutations in an antibiotic resistance gene. Mol Biol Evol 30:1779–87.

6. da Silva J, Coetzer M, Nedellec R, Pastore C, Mosier DE. 2010. Fitness epistasis and constraints on adaptation in a human immunodeficiency virus type 1 protein region. Genetics 185:293–303.

7. Lunzer M, Miller SP, Felsheim R, Dean AM. 2005. The biochemical architecture of an ancient adaptive landscape. Science 310:499–501.

8. Tan L, Serene S, Chao HX, Gore J. 2011. Hidden randomness between fitness landscapes limits reverse evolution. Phys Rev Lett 106:198102.

9. Gorter FA, Aarts MGM, Zwaan BJ, de Visser J. 2018. Local fitness landscapes predict yeast evolutionary dynamics in directionally changing environments. Genetics 208:307–22.

10. de Vos MG, Dawid A, Sunderlikova V, Tans SJ. 2015. Breaking evolutionary constraint with a tradeoff ratchet. Proc Natl Acad Sci U S A 112:14906–11.

11. Khan AI, Dinh DM, Schneider D, Lenski RE, Cooper TF. 2011. Negative epistasis between beneficial mutations in an evolving bacterial population. Science 332:1193–6.

12. Lalic J, Elena SF. 2015. The impact of high-order epistasis in the within-host fitness of a positive-sense plant RNA virus. J Evol Biol 28:2236–47.

13. Wistrand-Yuen E, Knopp M, Hjort K, Koskiniemi S, Berg OG, Andersson DI. 2018. Evolution of high-level resistance during low-level antibiotic exposure. Nat Commun 9:1599.

14. Salverda ML, Dellus E, Gorter FA, Debets AJ, van der Oost J, Hoekstra RF, Tawfik DS, de Visser JA. 2011. Initial mutations direct alternative pathways of protein evolution. PLoS Genet 7:e1001321.

15. Palmer AC, Toprak E, Baym M, Kim S, Veres A, Bershtein S, Kishony R. 2015. Delayed commitment to evolutionary fate in antibiotic resistance fitness landscapes. Nat Commun 6:7385.

16. Yang G, Anderson DW, Baier F, Dohmen E, Hong N, Carr PD, Kamerlin SCL, Jackson CJ, Bornberg-Bauer E, Tokuriki N. 2019. Higher-order epistasis shapes the fitness landscape of a xenobiotic-degrading enzyme. Nat Chem Biol 15:1120–28.

17. Mira PM, Meza JC, Nandipati A, Barlow M. 2015. Adaptive landscapes of resistance genes change as antibiotic concentrations change. Mol Biol Evol 32:2707–15.

18. Chou HH, Chiu HC, Delaney NF, Segre D, Marx CJ. 2011. Diminishing returns epistasis among beneficial mutations decelerates adaptation. Science 332:1190–2.

19. Bajic D, Vila JCC, Blount ZD, Sanchez A. 2018. On the deformability of an empirical fitness landscape by microbial evolution. Proc Natl Acad Sci U S A 115:11286–91.

20. Nicoloff H, Andersson DI. 2016. Indirect resistance to several classes of antibiotics in cocultures with resistant bacteria expressing antibiotic-modifying or -degrading enzymes. J Antimicrob Chemother 71:100–10.

21. Yurtsev EA, Chao HX, Datta MS, Artemova T, Gore J. 2013. Bacterial cheating drives the population dynamics of cooperative antibiotic resistance plasmids. Mol Syst Biol 9:683.

22. Yurtsev EA, Conwill A, Gore J. 2016. Oscillatory dynamics in a bacterial cross-protection mutualism. Proc Natl Acad Sci U S A 113:6236–41.

23. Butaite E, Baumgartner M, Wyder S, Kummerli R. 2017. Siderophore cheating and cheating resistance shape competition for iron in soil and freshwater Pseudomonas communities. Nat Commun 8:414.

24. Barlow M, Hall BG. 2002. Predicting evolutionary potential: in vitro evolution accurately reproduces natural evolution of the tem beta-lactamase. Genetics 160:823–32.

25. Frost I, Smith WPJ, Mitri S, San Millan A, Davit Y, Osborne JM, Pitt-Francis JM, MacLean RC, Foster KR. 2018. Cooperation, competition and antibiotic resistance in bacterial colonies. ISME J 12:1582–93.

26. Buijs J, Dofferhoff AS, Mouton JW, Wagenvoort JH, van der Meer JW. 2008. Concentration-dependency of beta-lactam-induced filament formation in Gram-negative bacteria. Clin Microbiol Infect 14:344–9.

27. Chow LKM, Ghaly TM, Gillings MR. 2021. A survey of sub-inhibitory concentrations of antibiotics in the environment. J Environ Sci 99:21–27.

28. Andersson DI, Hughes D. 2014. Microbiological effects of sublethal levels of antibiotics. Nat Rev Microbiol 12:465–78.

29. Gullberg E, Albrecht LM, Karlsson C, Sandegren L, Andersson DI. 2014. Selection of a multidrug resistance plasmid by sublethal levels of antibiotics and heavy metals. mBio 5:e01918–14.

30. Das SG, Direito SOL, Waclaw B, Allen RJ, Krug J. 2020. Predictable properties of fitness landscapes induced by adaptational tradeoffs. eLife 9:e55155.

31. Renggli S, Keck W, Jenal U, Ritz D. 2013. Role of autofluorescence in flow cytometric analysis of Escherichia coli treated with bactericidal antibiotics. J Bacteriol 195:4067–73.

32. Park SC, Neidhart J, Krug J. 2016. Greedy adaptive walks on a correlated fitness landscape. J Theor Biol 397:89–102.

33. de Visser JA, Park SC, Krug J. 2009. Exploring the effect of sex on empirical fitness landscapes. Am Nat 174 Suppl 1:S15–30.

34. Schenk MF, Zwart MP, Hwang S, Ruelens P, Severing E, Krug J, de Visser J. 2022. Population size mediates the contribution of high-rate and large-benefit mutations to parallel evolution. Nat Ecol Evol 6:439–47.

35. Sauvage E, Kerff F, Terrak M, Ayala JA, Charlier P. 2008. The penicillin-binding proteins: structure and role in peptidoglycan biosynthesis. FEMS Microbiol Rev 32:234–58.

36. Kummerer K. 2009. Antibiotics in the aquatic environment--a review--part I. Chemosphere 75:417–34.

37. Khan GA, Berglund B, Khan KM, Lindgren PE, Fick J. 2013. Occurrence and abundance of antibiotics and resistance genes in rivers, canal and near drug formulation facilities--a study in Pakistan. PLoS One 8:e62712.

38. Lindberg RH, Bjorklund K, Rendahl P, Johansson MI, Tysklind M, Andersson BA. 2007. Environmental risk assessment of antibiotics in the Swedish environment with emphasis on sewage treatment plants. Water Res 41:613–9.

39. Zuccato E, Calamari D, Natangelo M, Fanelli R. 2000. Presence of therapeutic drugs in the environment. Lancet 355:1789–90.

40. Zaborskyte G, Andersen JB, Kragh KN, Ciofu O. 2017. Real-time monitoring of nfxB mutant occurrence and dynamics in Pseudomonas aeruginosa biofilm exposed to subinhibitory concentrations of ciprofloxacin. Antimicrob Agents Chemother 61.

41. Westhoff S, van Leeuwe TM, Qachach O, Zhang Z, van Wezel GP, Rozen DE. 2017. The evolution of no-cost resistance at sub-MIC concentrations of streptomycin in Streptomyces coelicolor. ISME J 11:1168–78.

42. Jorgensen KM, Wassermann T, Jensen PO, Hengzuang W, Molin S, Hoiby N, Ciofu O. 2013. Sublethal ciprofloxacin treatment leads to rapid development of high-level ciprofloxacin resistance during long-term experimental evolution of Pseudomonas aeruginosa. Antimicrob Agents Chemother 57:4215–21.

43. Matange N, Hegde S, Bodkhe S. 2019. Adaptation through lifestyle switching sculpts the fitness landscape of evolving populations: implications for the selection of drug-resistant bacteria at low drug pressures. Genetics 211:1029–44.

44. Payne JL, Menardo F, Trauner A, Borrell S, Gygli SM, Loiseau C, Gagneux S, Hall AR. 2019. Transition bias influences the evolution of antibiotic resistance in Mycobacterium tuberculosis. PLoS Biol 17:e3000265.

45. Ruelens P, de Visser JA. 2021. Choice of beta-lactam resistance pathway depends critically on initial antibiotic concentration. Antimicrob Agents Chemother 65:e0047121.

46. Barbosa C, Romhild R, Rosenstiel P, Schulenburg H. 2019. Evolutionary stability of collateral sensitivity to antibiotics in the model pathogen Pseudomonas aeruginosa. eLife 8:e51481.

47. Roemhild R, Linkevicius M, Andersson DI. 2020. Molecular mechanisms of collateral sensitivity to the antibiotic nitrofurantoin. PLoS Biol 18:e3000612.

48. Cristovao F, Alonso CA, Igrejas G, Sousa M, Silva V, Pereira JE, Lozano C, Cortes-Cortes G, Torres C, Poeta P. 2017. Clonal diversity of extended-spectrum beta-lactamase producing Escherichia coli isolates in fecal samples of wild animals. FEMS Microbiol Lett 364.

49. Pai H, Lyu S, Lee JH, Kim J, Kwon Y, Kim JW, Choe KW. 1999. Survey of extended-spectrum beta-lactamases in clinical isolates of Escherichia coli and Klebsiella pneumoniae: prevalence of TEM-52 in Korea. J Clin Microbiol 37:1758–63.

50. Ojkic N, Serbanescu D, Banerjee S. 2022. Antibiotic resistance via bacterial cell shape-shifting. mBio 13:e0065922.

51. Melnyk AH, Wong A, Kassen R. 2015. The fitness costs of antibiotic resistance mutations. Evol Appl 8:273–83.

52. Johnson JB, Omland KS. 2004. Model selection in ecology and evolution. Trends Ecol Evol 19:101–8.

